# Insect wings arose with a genetic circuit that extends the useful range of a BMP morphogen

**DOI:** 10.1101/2025.02.07.637038

**Authors:** Anqi Huang, Luca Cocconi, Ben Nicholls-Mindlin, Cyrille Alexandre, Guillaume Salbreux, Jean-Paul Vincent

## Abstract

Morphogens are produced by a subset of cells to trigger a signalling gradient that provides positional information to surrounding tissues. At increasing distances from the source, the dwindling number of morphogen molecules is expected to constrain the useful range of morphogen gradients. We have identified a genetic circuit that counteracts this limitation in developing wings of *Drosophila* by boosting BMP signalling at the distal end of the gradient without amplifying the signal near the source. This circuit involves Brinker, a transcription factor that represses BMP target genes while itself being repressed by BMP signalling. We suggest that temporal averaging inherent to the production of the inverse Brk gradient contributes to the enhancement of the positional information far from the Dpp source. Despite being a core component of BMP signalling in flies, Brinker is exclusively found in insects, likely in all insect species. Genomic analysis across a wide range of insects and gene expression analysis in limb primordia of the apterygote *Thermobia domestica* suggests that Brinker is an insect-specific innovation that was subsequently wired into the BMP signalling network in pterygotes, perhaps to enable wing development.

## Main Text

### Graded Dpp signalling triggers a reverse transcriptional gradient of *brinker*

The appearance of wings was a momentous event in evolution, setting the stage for the diversification of the most successful group of animals on the planet (*1,2,3*). The wings of holometabolous insects develop from imaginal discs, epithelial pockets that grow during larval stages (*4,5*). Like many developing tissues, they rely on morphogens for positional information (*6,7,8*). In *Drosophila melanogaster* (*Dm*), the anterior-posterior (A-P) axis of wing imaginal discs is patterned by a gradient of Dpp, a BMP homolog, which is maintained until the end of the third instar when the primordium is about 100 cell diameters wide (*9,10,11,12,13*). It is expected that in such a large developmental field, the dwindling number of morphogen molecules far from the source would constrain the useful range of morphogen gradients (*14,15,16,17*). This limitation is often overcome in diverse developmental systems by an independent reverse morphogen gradient (Fig. 1A, B) (*18*). For example, in the vertebrate neural tube, opposing gradients originating from sources specified by tissue-extrinsic landmarks (the ventral notochord and the dorsal ectoderm) ensure positional precision across the dorso-ventral axis (*19*). Imaginal discs, however, lack these independent landmarks as they develop semi-autonomously from the rest of the animal. Instead, wing imaginal disc’s morphogens are produced from stripes of cells established during embryogenesis before disc primordia invaginate to form imaginal discs. Thus, the Dpp morphogen is produced from the A-P boundary, which derives from the embryonic parasegment boundary that cuts across the second thoracic segment. During the 4-day period of larval development, the Dpp signalling gradient can be detected with antibodies against phosphorylated Mad, a key transducer of the pathway (Fig. 1C-E). There is no evidence for an independent reverse morphogen gradient along the A-P axis. The transcription factor encoded by *brinker* (*brk*) does form a reverse gradient but it is thought to be entirely determined by pMad-mediated repression (*20,21,22*). Indeed, Brk expression becomes uniformly high in response to acute removal of Dpp by optogenetic inactivation, confirming that the Brinker gradient depends on the Dpp gradient (Fig.1 F-H). We created an authentic *brk* transcriptional reporter by knocking DNA encoding mCherry at *brk*’s endogenous translation initiation codon (*brk[KO; mCherry]*) and this was found to be expressed in a smooth reverse gradient, confirming that the reverse Brk gradient is transcriptionally generated, in response to the pMad gradient (Fig. 1I). Aberrant Brk expression results in patterning defects and abnormal growth, hence leading to abnormal wing shape (Fig. S2), indicating that the Brk gradient is essential for normal development. It is thought that, together, the pMad and Brk gradients control the expression of targets such as *spalt major* (*salm*) and *optomotor-blind* (*omb*), which in turn specify the position of patterning elements such as wing veins (*23,24,25*). Therefore, two opposite gradients provide positional information along the A-P axis although the benefit in terms of positional information is unclear since one gradient depends on the other.

**Figure. 1.**
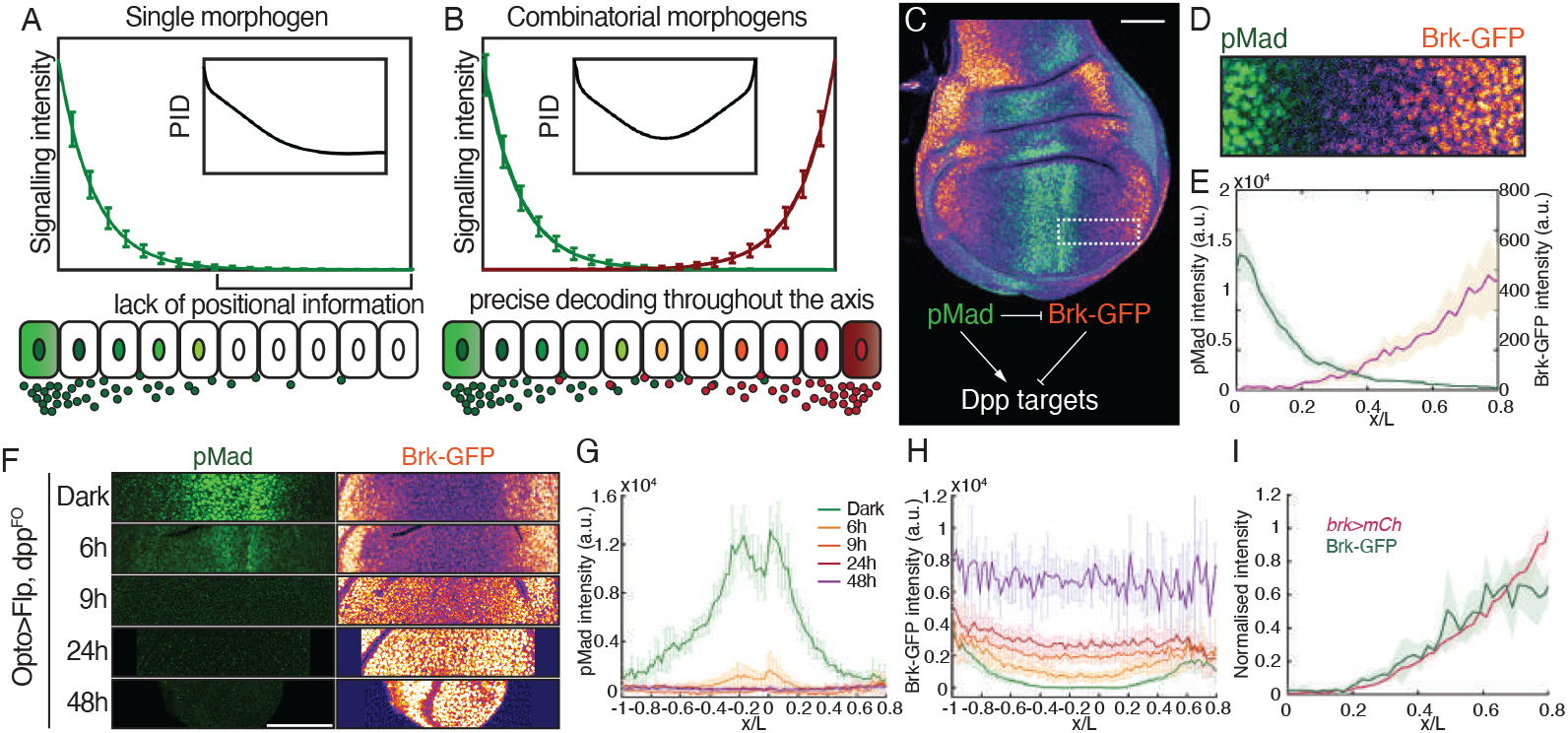
Graded Dpp signalling is translated into a reverse transcriptional gradient of *brk*. (**A**) Decoding morphogen signals is particularly challenging far from the source, where stochastic molecular interactions obscure reliable readouts, leading to reduced positional information density (PID). (**B**) Opposing morphogen gradients can provide accurate positional information throughout the patterning axis. Synthetic data for (A-B) was generated by adding normal noise to exponential gradients with PID calculated as in Methods, section 1.11. (**C**-**E**) *Drosophila* wing imaginal disc expressing Brk-GFP from a knock-in allele stained with anti-pMad. The area marked by a dashed rectangle in (C) is shown at higher magnification in (D). Scale bar: 50 µm. Mean profiles of pMad immunoreactivity and Brk-GFP fluorescence in the ventral-posterior compartment of 9 wing discs at 120 hours after egg lay (AEL), plotted against relative A-P position (E). Nuclei were segmented and binned every 2% *x/L*. (**F**-**H**) Brk-GFP wing discs stained with anti-pMad after light-induced excision of *dpp*. Scale bar: 25 µm. The mean pMad (G) and Brk (H) profiles are shown as functions of relative A-P position at different time points after *dpp* inactivation. (**I**) Normalised intensity of fluorescence from a knock-in *brk>mCh* transcriptional reporter (*brk* null mutant) and Brk-GFP (fully functional), present on homologous chromosomes, plotted against relative A-P position.

### Brk boosts positional information by modulating the pMad gradient

Although the Brk gradient is clearly specified by Dpp signalling activity, as we now show, it is not merely a passive readout of the pMad gradient. This is readily seen with a conditional allele (*brk[KO; FRT-BrkV5-FRT]*) that can be inactivated by Flp recombinase in specific tissue domains (e.g. in the dorsal compartment), leaving the complementary domain (e.g. the ventral compartment) as control. In the absence of functional Brk (upper compartment in Fig. 2A), the *brk* reporter no longer forms a smooth gradient and instead is expressed in a step-like manner, low near the Dpp source and high beyond 40% of the A-P axis. Therefore, Brk function is required for *brk*’s transcriptional gradient. Moreover, in the *brk* mutant domain, Dpp signalling, as measured by pMad, was seen to drop more sharply than in the control compartment, suggesting that Brk positively feeds back on the pMad gradient. In the absence of Brk, no such feedback takes place, and positional information density plots (see Methods, section 1.11) show quantitatively that the positional information available from the pMad gradient or the nominal *brk* transcription gradient is reduced at the distal end of the Dpp gradient (compare red mutant to blue control curves in Fig. 2E, F). We also calculated, for each gradient, the ‘expected *a posteriori* distribution’, where higher intensity along the diagonal indicates greater positional precision (see Methods, section 1.10) (*26, 27, 28*). These maps show that beyond position 0.4 of the control region, the Brk gradient provides more reliable positional information than the pMad gradient (Fig. 2G, H). In the absence of Brk activity, neither gradient is capable of accurately indicating position in this region (Fig. 2I, J). Therefore, positive modulation of pMad by Brk contributes to positional precision in the distal region of the Dpp gradient. When the information content from both gradients is combined, the beneficial role of Brk becomes even more evident (Fig. 2K, L).

**Figure. 2.**
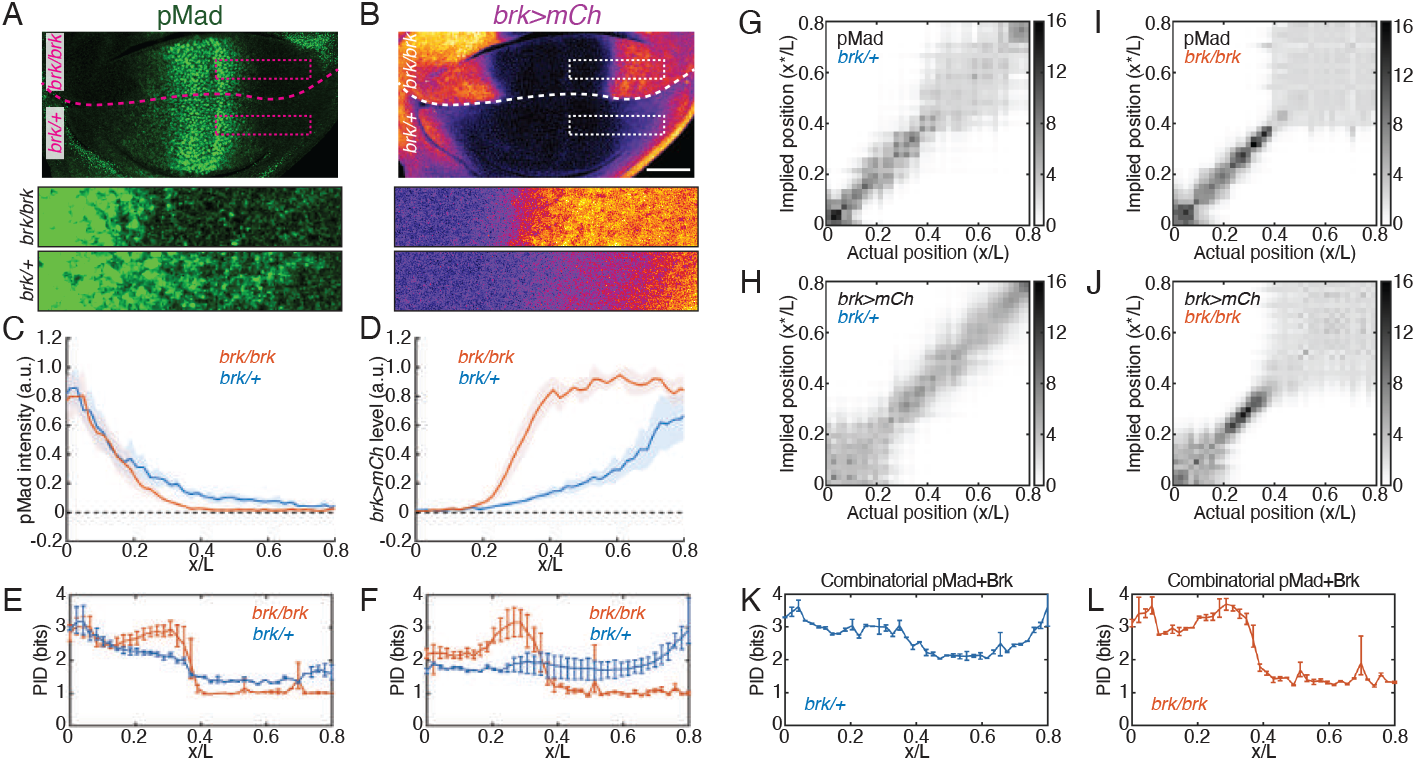
Impact of *brk* mutation on pMad and *brk* transcription profiles. (**A**-**B**) Wing discs harbouring the *brk>mCh* transcriptional reporter (*brk* null mutant) and a *brk* conditional allele on the homologous chromosome. *ap>Gal4* drives Flp to inactivate the conditional allele and make the dorsal compartment *brk* mutant. The profiles of pMad and *brk>mCh* were quantified in both the dorsal and ventral compartments, with the highlighted areas shown at higher magnification underneath. In the magnified pMad images, the maximum displayed intensity was adjusted to a lower threshold to enhance contrast and improve visualization of differences in pixel intensity at lower values. Scale bar: 50 µm. (**C**-**D**) Mean profiles of pMad (C) and *brk>mCh* (D) plotted against *x/L*. The dashed lines indicate the background level. (**E**-**F**) Positional information density of the pMad (E) and *brk* (F) profiles in control and *brk* mutant domains. The prior distribution is defined over the spatial range of [0, 0.8], resulting in a PID of approximately 1 bit in the distal half of the mutant gradients, where the average pMad and *brk* profiles plateau beyond *x/L* = 0.4. (**G**-**J**) Expected *a posteriori* probability distributions of pMad (G, I) and *brk* (H, J) profiles in the control (G, H) or *brk* mutant (I, J) domains. (**K**-**L**) Positional information density derived from the combinatorial gradients of pMad and nominal *brk* in control (K) and *brk* mutant (L) conditions.

We next address how Brk modulates the pMad gradient. The level of pMad is determined by the rates of phosphorylation and dephosphorylation. Mad phosphorylation is achieved by Tkv, a conserved type I BMP receptor, and this is counteracted by Daughters against Dpp (Dad), a conserved inhibitory Smad (*29,30,31*). Therefore, Brk could boost Mad phosphorylation by upregulating Tkv or downregulating Dad. This was tested by assessing the effect of *brk* loss of function on the expression of the transcriptional reporters *tkv>lacZ* and *dad>lacZ*. Compared to the levels seen in wildtype tissue, expression of *dad>lacZ* was enhanced and that of *tkv>lacZ* was reduced in the *brk* mutant domain despite unchanged Dpp morphogen production (Fig. 3A-G). Tkv protein was also reduced, as indicated by the activity of a fusion knock-in allele, *tkv-GFP* (Fig. 3E). Therefore, Brk normally represses *dad* and promotes *tkv*. To assess a potential effect of Brk on pMad dephosphorylation, we quantified the time course of pMad levels in Brk overexpressing and control tissues after acute Tkv inhibition with LDN-193189, a small molecule that prevents type I BMP receptors from phosphorylating Smads. No difference in the decay rate was seen, suggesting that Brk does not modulate pMad dephosphorylation (Fig. S3). Overall, the above results suggest that a gene regulatory network (GRN) involving feedback regulation of *tkv* and *dad* transcription could account for the enhancement of Mad phosphorylation by Brk (Fig. 3H).

**Figure. 3.**
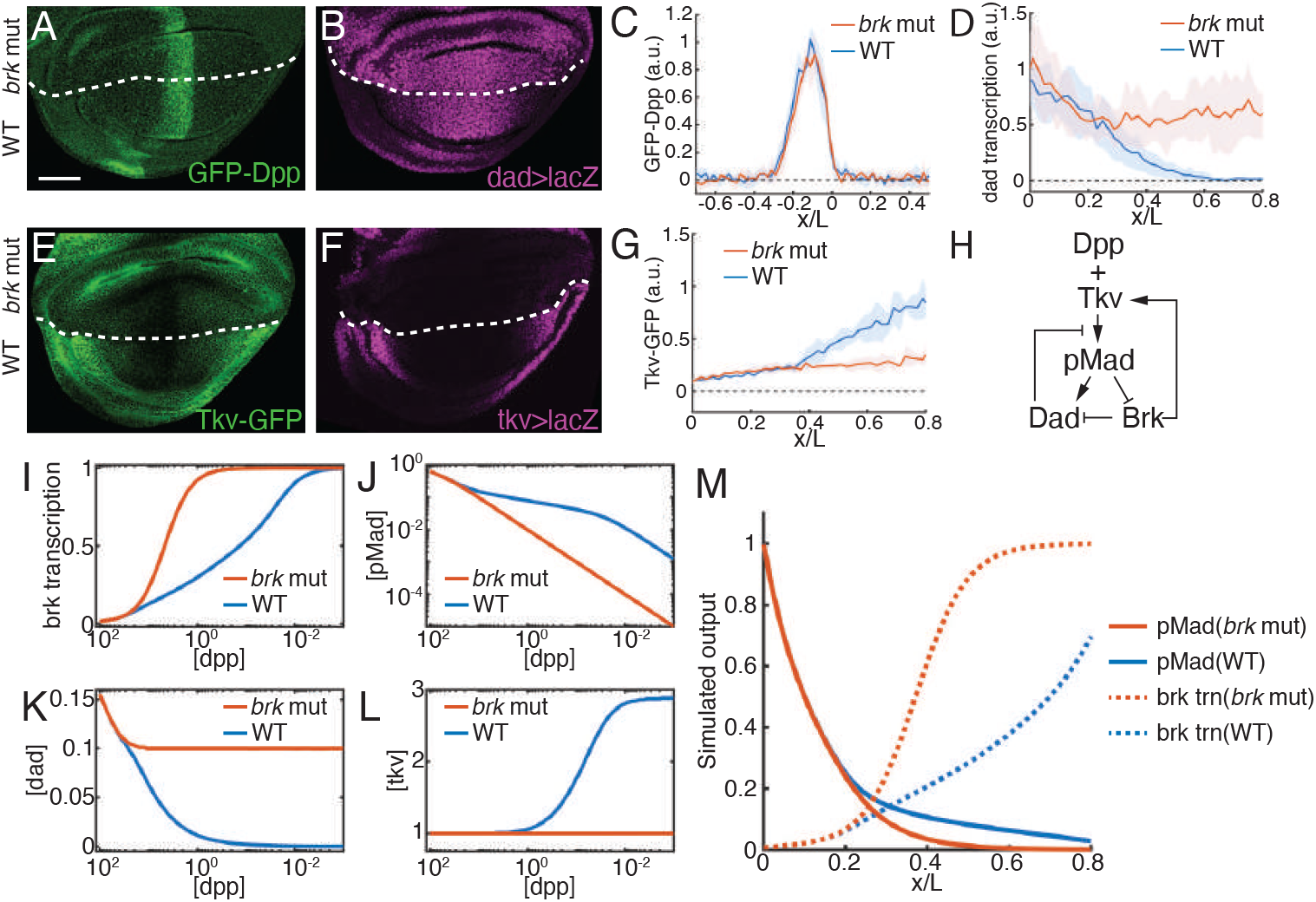
Brk-mediated transcription regulation of the Type I receptor Tkv and the inhibitory Smad, Dad. (**A**-**G**) Wing discs lacking *brk* activity in the dorsal compartment, expressing GFP-Dpp from a knock-in allele (A), *β*-galactosidase from a *dad>lacZ* enhancer trap (B), a Tkv-GFP fusion from a knock-in allele (E), or *β*-galactosidase from a *tkv>lacZ* enhancer trap (F). Mean spatial profiles of GFP-Dpp (C), *dad>lacZ* (D), or Tkv-GFP (G) in the wild type (WT, blue) and *brk* mutant (*brk* mut, red) compartments. Scale bar: 50 µm. (**H**) Regulatory interactions between Brk and Dpp signalling components. (**I**-**L**) Simulated steady-state values of signal transduction components (pMad, *brk, dad*, and *tkv*), plotted against extracellular Dpp concentration in the presence or absence of Brk activity. These plots represent numerical solutions of the model’s cell-autonomous module, using the fitted parameters listed in Table S5 in the Supplementary Text. Reference units for the concentrations of different species are discussed in the Supplementary Text; for instance, extracellular pMad concentration is normalised to its value at the A-P boundary of wildtype discs. (**M**) Simulated spatial profiles of pMad and *brk* transcription in wild type and *brk* mutant conditions.

For formal evaluation of the Brk feedback circuit, we developed a mathematical model consisting of two core modules (see Supplementary Text). Module 1 accounts for extracellular Dpp transport via a diffusion equation that includes receptor binding dynamics and a localized source term. At steady-state and in the posterior compartment, the extracellular concentration of free Dpp, denoted [*dpp*], satisfies

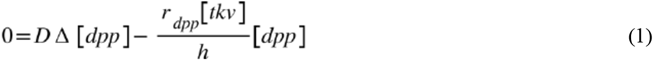

with *D* the ligand diffusivity, *r*_dpp_ an effective degradation rate per unit of receptor volume concentration in the intercellular space, *h* the width of the intercellular space and [*tkv*] the membrane concentration of free receptors. Module 2 describes cell-autonomous intracellular signalling dynamics, whereby Dpp binding to Tkv induces Mad phosphorylation, which is competitively inhibited by Dad. The rate of pMad dephosphorylation is assumed constant.

Accordingly, the steady-state intracellular concentration of pMad, denoted [*pMad*], satisfies

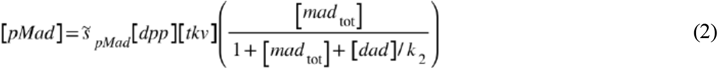

with 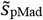 a parameter related to the ratio of Mad phosphorylation and dephosphorylation rates, [*mad*_tot_] and [*dad*] the concentrations of total Mad and Dad, *k*_1_ and *k*_2_ the binding affinities for Tkv of Mad and Dad, respectively. This module also incorporates Brk’s activation of *tkv* and repression of *dad*, along with the repression of *brk* by pMad (Supplementary Text, section 2.1). Positive and negative regulatory interactions were formalised via non-linear Hill functions with parameters fitted directly to empirical input-output response curves. Other parameters that are not currently available in the literature, e.g. the relative concentrations of intracellular species, were also extracted from experimental measurements (Supplementary Text, section 2.2). Solving module 2 equations using the fitted parameters allowed us to derive the steady-state values of the intracellular components as a function of extracellular Dpp (Fig. 3I-L). We then drew on these response curves to solve the stationary signal profiles predicted by the diffusion-degradation equation in module 1, producing a set of spatial profiles, which recapitulated the gene expression patterns observed in both wildtype and *brk* mutant wing discs (Fig. 3M). This analysis showed that the Brk circuit is sufficient to extend the pMad gradient and to shape the Brk gradient.

Next, we assessed the relative importance of the two feedback branches *in vivo* and *in silico*. For an *in vivo* assessment, we used the Gal4 system to reintroduce these feedback branches individually in *brk* mutant tissue, which inherently lacks both branches. To test the relevance of the *dad* branch, Gal4 expressed from a *brk[KO; Gal4]* allele was used to drive simultaneously, in the *brk* domain, an RNAi transgene against *dad* and a UAS*>*Flp transgene to inactivate a *brk* conditional allele, *brk[KO; FRT-Brk-FRT]*, which also serves as a *brk* transcriptional reporter (see Materials and Methods, section 1.2.1). In the resulting larvae, the pMad gradient was extended and the reverse transcriptional *brk* gradient was partially restored (Fig. 4A-B), suggesting that feedback regulation of Dad expression by Brk could suffice for the establishment of a gradual Brk expression profile in the distal region of the Dpp gradient. No such effect was seen with Tkv overexpression (Fig. 4A, C), suggesting that the Dad repression branch could be more relevant than that involving Tkv upregulation. We then tested our model’s response to analogous *in silico* manipulations, using previously fitted parameters. As *in vivo*, virtual Dad downregulation extended Dpp signalling and smoothed the Brk gradient, while boosting virtual Tkv had minimal effect (Fig. 4B’, C’). Further experimental and simulation tests, including replacing endogenous Tkv with uniformly expressed Tkv or introducing a *dad* mutation, yielded corroborative results (Supplementary text, section 2.4). Specifically, uniform Tkv had little impact on the reverse Brk gradient, suggesting that the Tkv feedback branch is not critical for its formation (Fig. S9E-F). In contrast, removal of Dad activity by mutation allowed for precise decoding in the gradient’s distal region, underscoring the importance of Dad repression in establishing the reverse Brk gradient. In a *dad* mutant, since Dad is a feedback inhibitor of signalling, global loss of Dad activity would be expected to also boost signalling in the proximal region of the gradient. This was not observed, although a departure from the wildtype pMad gradient was observed around 40% tissue length in *dad* mutant imaginal discs (Fig. S10; see also Supplementary Text 2.4.3). This suggests the existence of an additional feedback mechanism that dampens signalling near the Dpp source. Nevertheless, our model accurately represents the circuit that generates the reverse Brk gradient far from the Dpp source and we conclude that repression of *dad* by Brk is essential for decoding the Dpp gradient’s distal end.

**Figure. 4.**
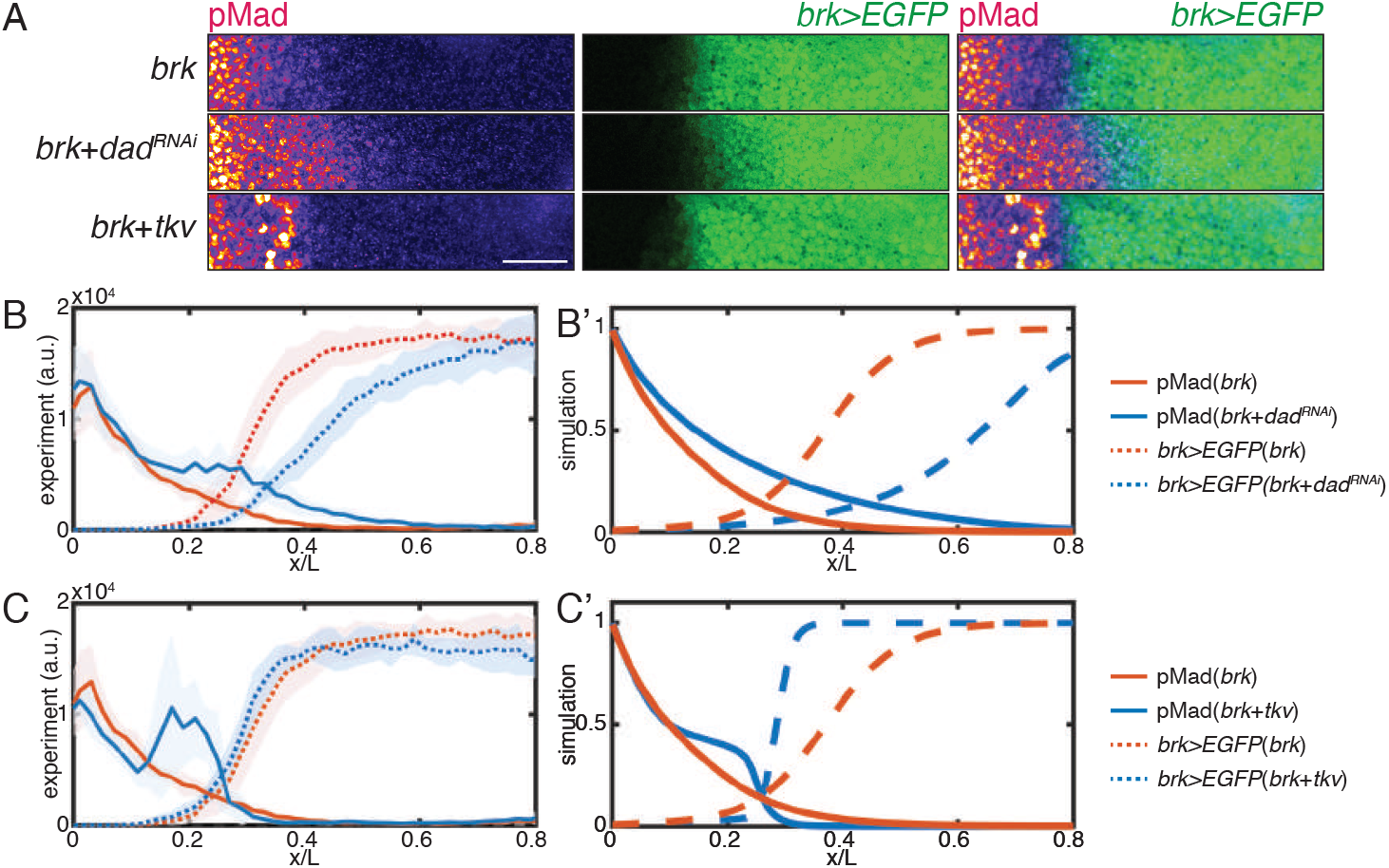
Effect of the two feedback branches *in vivo* and *in silico*. (**A**) Top panel shows pMad immunoreactivity and *brk>EGFP* reporter activity in homozygous *brk* mutant imaginal discs (*brk>EGFP*/*brk>Gal4*; see detailed genotypes in Methods). The other panels show the same genotype with, in addition, *UAS>dad*^*RNAi*^ or *UAS>tkv*. Scale bar: 25 µm. (**B**-**C**) Mean spatial profiles of pMad (solid lines) and *brk>EGFP* activity (dashed lines) measured in *brk* mutant discs with *UAS>dad*^*RNAi*^ (B) or *UAS>tkv* (C) in blue, compared to the *brk* mutant alone in red. Simulated counterparts of the profiles are shown in (B’-C’).

### The *brinker* gene is an Insecta innovation

The BMP signalling network is highly conserved across the animal kingdom and plays an essential role in patterning a wide range of tissues (*32, 33, 34, 35*). Yet, homologs of Brk, an essential component of BMP signalling in *Drosophila*, have not been reported outside insects (*36, 37*). To investigate the evolutionary origin of Brk, we conducted tblastn searches using the *Drosophila* Brk (Brk^Dm^) protein sequence against translated representative genomes of the class Insecta. This search led to the identification of Brk homologs in all 123 species examined, with high alignment (similarity*>*80%) within the Brk DNA binding domain (Data S1). These homologs vary in terms of the presence of recognisable co-repressor recruitment domains (*38,39*), as well as the number and conservation of the DNA-binding domains (Fig. 5A, Table S1) (*40,41,42*). Brk from holometabolous insects have at least one recognisable co-repressor recruitment domain, while such domains could not be identified in more basal hemimetabolous and ametabolous insects (*43*). Instead, species of the latter groups tend to have additional (up to three) DNA-binding domains, potentially enhancing specificity and efficiency of DNA recognition and target search (*44, 45*). Based on alignment levels between different DNA binding domains within a protein and between different species, the triple DNA binding domain version of the Brk protein appears to be the more ancient form (Fig. S4A). During evolution, this version seems to have lost the third domain (as in Orthoptera and Hymenoptera), the second domain (as in Coleoptera), or both domains (as in Lepidoptera and Diptera). Brk homologs could also be identified in non-insect hexapods, although they appear to only retain homology in the DNA binding domain and differed from insect Brk in their genomic organisation (Fig. S4C-E). Notably, no Brk homolog could be identified outside hexapods, even in other arthropods such as crustaceans (see Methods section 1.12). In conclusion, our analysis suggests that Brk is a DNA-binding protein that was only maintained in hexapods during evolution.

**Figure. 5.**
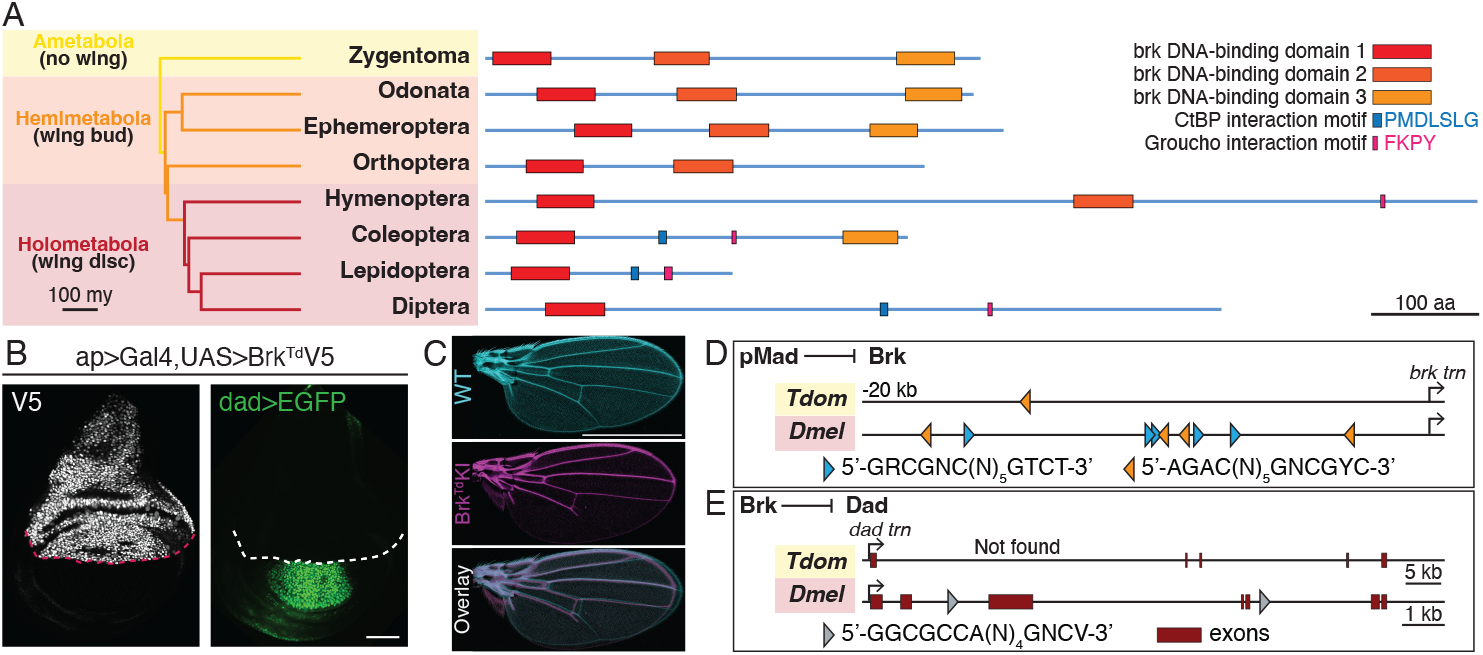
The *brk* gene is an evolutionary innovation of insects. (**A**) Functional motifs of Brk proteins for multiple representative orders within the class Insecta. (**B**) *Drosophila* wing disc carrying the *dad>EGFP* reporter and expressing V5-tagged *Thermobia* Brk from a UAS-transgene in the dorsal compartment. Dashed lines indicate the D-V boundary. Scale bar: 50 µm. (**C**) *Drosophila* wing expressing *Thermobia* Brk instead of *Drosophila* Brk from the endogenous locus (Brk^Td^KI) compared to control wing from wild type OregonR strain (WT). Scale bar: 1 mm. (**D**) pMad-Med-Shn silencer motifs within 20kb upstream of the *brk* TSS in *Drosophila* and *Thermobia*. Blue and orange colours indicate 5’ or 3’ orientation. (**E**) Brk-binding sites (grey triangles) located within introns of *Drosophila dad*. No such sites were identified within the *Thermobia dad* gene.

While sequence-specific DNA-binding is a central feature of Brk, one cannot predict from sequence alone whether Brk’s original function was to repress the transcription of BMP target genes. To explore this, we turned to *Thermobia domestica* (*Td*), a member of the basal insect order Zygentoma (Fig. 5A). This allowed us to determine if Brk’s role as a transcriptional repressor has been conserved across large evolutionary distances and within distant wing-less insects (apterygotes) (*46,47,48*). We evaluated Brk^Td^’s activity in *Drosophila* by overexpressing V5-tagged Brk^Td^ in wing imaginal discs. This led to marked suppression of Dpp targets *salm, omb* and *dad* (Fig. 5B, Fig. S5A-E), indicative of repressor function, despite the absence of recognisable co-repressor recruitment domains. It is remarkable that known Brk^Dm^ target genes are repressed by Brk^Td^ in *Drosophila*, suggesting that Brk’s function has been maintained across the insect class. Functional conservation was further tested by replacing, in *Drosophila*, the endogenous Brk coding sequence with that of Brk^Td^ to generate *brk[KO; Brk*^*Td*^] animals. Null *brk* mutations (e.g. *brk[KO]*) are lethal in *Drosophila* because of excess expression of Dpp signalling target genes in various tissues (*49*). In contrast, a small proportion of homozygous *brk[KO; Brk*^*Td*^] flies survived with near normal wings (Fig. 5C, Fig. S5F-H). We conclude that the molecular function of Brk has remained largely unchanged during insect evolution, most likely to dampen transcriptional targets of Dpp signalling (*50*).

For the Brk module to extend the read-out precision of the Dpp gradient, two conditions must be fulfilled (Fig. 3H). One is that Brk represses *dad* expression, as it does in *Drosophila* (see below for *Thermobia*). The other, which we now address, is that *brk* itself be repressed by Dpp signalling. In *Drosophila*, transcriptional repression of *brk* by Dpp signalling is mediated by pMad silencer motifs, which recruit a tripartite complex comprising pMad, Medea and the co-repressor Schnurri (Shn) (*51,52,53,54*). Remarkably, the vertebrate homologs Smad1, Smad4, and Shn also form a complex that recognizes the same DNA sequence (*55*). Therefore, pMad silencer sites can readily be identified by sequence homology (Fig. 5D) and their presence can be used as an indication that nearby genes could be regulated by Dpp signalling (*56*). Nine pMad silencer sites can be found within 20kb upstream of the *brk*’s transcription start site (TSS) in *Drosophila*. The number and topology of these sites are relatively conserved across closely related species, an indication of their functional relevance. In more distant holometabolous species, the sites are still present but in reduced numbers (down to four in Hymenoptera). This number is further reduced in hemimetabola, with 2-3 sites found 20kb upstream of *brk*’s TSS (Table S2, Fig. S5I). Previous studies of *Gryllus* (an Orthopteran of the hemimetabola group) have shown that *brk* transcription increases in response to *mad* RNAi at the nymph stage (*57*), consistent with repression by BMP signalling. It is likely therefore that the pMad silencer sites found upstream of *brk* are functional in this species, and by extension in other hemimetabola. Therefore, our analysis so far suggests that the precision-enhancing Brk module is operational in pterygote insects. We now address whether this is likely to be true in apterygotes.

To assess the connectivity of the Brk module in apterygotes, we began by exploring whether Dpp signalling is likely to repress *brk* expression in this subclass of insects. Only one pMad silencer motif could be identified upstream of *brk*, about 14kb upstream of the TSS, in the apterygote *Thermobia domestica* (Fig. 5D). It is doubtful whether a single element located so far from the TSS would suffice to mediate repression by pMad. However, one cannot exclude the presence of other silencer motifs further than 20kb from the TSS. Indeed, it is problematic to infer repression by pMad from sequence information alone. To address this experimentally in *Thermobia domestica*, we established a colony in our laboratory and began to characterise the relationship between *brk* expression and BMP signalling in developing appendages of this species (Fig. 6A-G). Dpp was found to be expressed at the ectodermal tip, while uniform low-level expression could be detected throughout the mesoderm (Fig. S6A). Immunofluorescence staining with anti-pMad, and HCR to detect *dad* transcripts, two indicators of Dpp signalling, confirmed that localised Dpp expression leads to a distal-to-proximal gradient of signalling activity. Yet, *brk* was expressed at a uniform low level (see Fig. S6B for probe validation), not in a reverse gradient, suggesting that *brk* expression is not modulated by Dpp signalling in *Thermobia*. We tested this conclusion by assessing *brk* expression in *Thermobia* after 1.5 hr treatment with LDN-193189 to prevent Dpp signalling (Fig. 6H-K). As expected, pMad immunoreactivity and the *dad* HCR signal were strongly reduced, indicating that Dpp signalling was impaired. Yet, *brk* expression remained unchanged. We conclude that, in *Thermobia domestica*, Brk expression is not significantly regulated by Dpp signalling. Therefore, a key requirement of the Brk feedback module is not fulfilled in this species. The other requirement, inhibition of *dad* expression by Brk cannot be rigorously tested for lack of genetic means of knocking down/out Brk activity. We note however the absence of recognisable Brk binding sites in the intronic region of *Thermobia*’s *dad* while such sites are abundant in the intronic region of pterygotes (Fig. 5E; Fig. S5J). We conclude that, even though Brk^Td^ is a transcription repressor, it is unlikely to regulate *dad* expression in *Thermobia domestica*. Overall, the above analysis, suggests that the precision-enhancing Brk module is unlikely to be operational in this wing-less species (Fig. 5L).

**Figure. 6.**
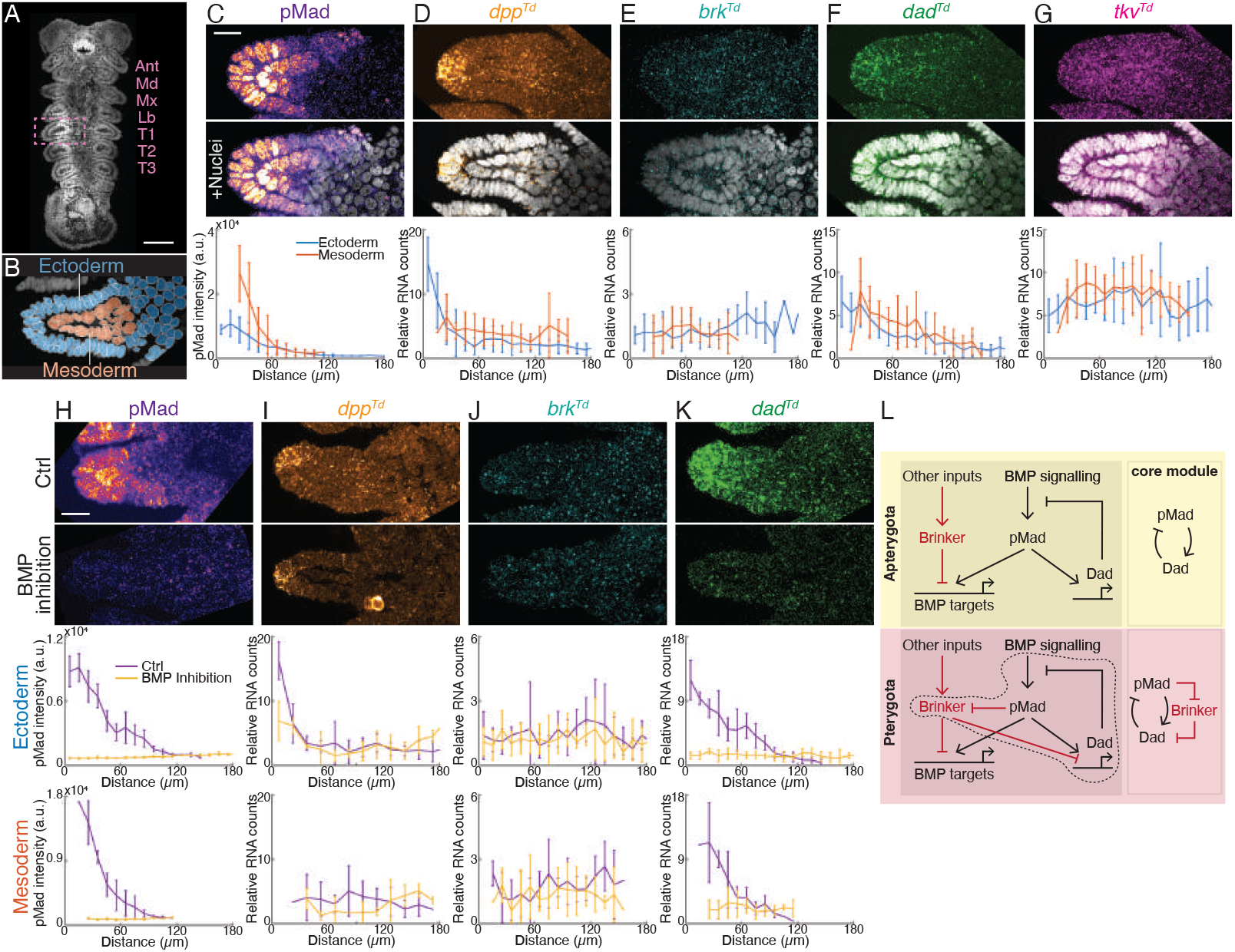
The Brk network is incomplete in *Thermobia* limbs. (**A**) Midsagittal view of a *Thermobia* embryo during early limb extension, with nuclei labelled. The dashed rectangle highlights the limb of the first thoracic segment. Abbreviations: Ant, antenna; Md, mandible; Mx, maxilla; Lb, labium; T1-T3, first to third thoracic segments. Scale bar: 100 µm. (**B**) Segmentation of nuclei within the T1 limb primordium, with blue and red indicating ectodermal and mesodermal nuclei, respectively. (**C**-**G**) T1 limb primordia stained with anti-pMad (C) or with transcripts detected by *in situ* hybridization for *dpp*^*Td*^ (D), *brk*^*Td*^ (E), *dad*^*Td*^ (F), or *tkv*^*Td*^ (G). The same images overlaid with nuclear staining are shown underneath. The plots show mean pMad intensity (C) or mean transcript counts per cell (D-G) plotted against distance from the limb distal end. Scale bar: 25 µm. (**H**-**K**) Limb primordia from control (top) or LDN-193189-treated (bottom) *Thermobia* embryos stained for pMad (H) or with transcripts detected by *in situ* hybridization for various genes (I-K). Plots show spatial profiles of pMad intensity and transcript counts per cell in ectodermal and mesodermal tissues. Scale bar: 25 µm. (**L**) Regulatory interactions of Brk with BMP signalling components in apterygote and pterygote insects. Evolutionary innovations are marked in red: Brinker itself in all insects (top and bottom) and linkage to the Dad feedback loop in pterygotes (bottom). The dashed line encloses the part of the network that is essential for signalling gradient extension. Core network modules are illustrated on the right.

## Discussion

In this study, we have identified and characterised a genetic circuit that extends the spatial range over which BMP signalling can be decoded. This module acts on top of the highly conserved negative feedback mechanism whereby pMad activates expression of an inhibitory Smad (Dad in *Drosophila*) (*58,59,60,61*). Although formally, the Brk circuit acts as an additional negative feedback module, its primary effect is to enhance BMP signalling specifically in the region of the gradient where signalling is particularly low. Thus, at the distal end of the gradient, Dad-mediated negative feedback is alleviated by Brk-mediated repression, lifting Dpp signalling solely in this low-signal region. Consequently, the Brk network acts as an intensity-modulated signal amplifier that sensitises cells to Dpp only at low concentrations. This mechanism is reminiscent of an electronic logarithmic amplifier, which increases the gain as the input signal decreases. Examples of logarithmic sensing in biology can be found in sensory perception and bacterial aerotaxis and have been predicted in biochemical circuits (*62,63,64,65*). Despite the expectation that amplification would increase noise along with the signal, the Dpp gradient still provides sufficient positional information across the A-P axis, as evidenced by the high positional information density in the reverse Brk gradient. We propose that noise is mitigated through temporal integration: while pMad provides a relatively instantaneous readout of the local extracellular ligand concentration (*t*_1*/*2_=14 min, Fig. S3), Brk protein, with its longer half-life (*t*_1*/*2_=10 hrs), achieves some temporal signal integration. Extension of the range over which the pMad gradient can provide positional information could be particularly relevant to insects, which set aside their wing primordia during their semi-aquatic larval stage (*66*). Since insect wings must develop without positional cues from the rest of the organism and are relatively large, they must maximize the positional information that can be achieved from individual morphogen gradients. Remarkably, in *Drosophila*, the role of the Brk module in signalling extension is most prominent in wing disc development compared to other developmental processes such as embryonic and germline patterning (*49,67*). This prominence is likely due to the proximity of Brk wing enhancers to pMad silencer sites, making Brk expression highly sensitive to pMad repression (*20,68*). It is worth noting that the Brk network seems to have become established concurrently with the emergence of wings. It is plausible therefore that this network’s configuration facilitated precise wing patterning in flying insects. However, since Brk expression is modulated by Dpp signalling in a variety of tissues in modern insects, we cannot exclude the possibility that the Brk module enabled innovation elsewhere, albeit still specifically in winged insects. We suggest that the Brk module evolved in two stages in an ancestor that already had a functional BMP signalling pathway, including inhibitory Smad (Dad)-mediated negative feedback. First, a novel transcription factor (Brk) evolved, perhaps to dampen BMP targets in an as-of-yet unidentified tissue. Subsequent emergence of pMad regulatory elements in the *brk* gene and the appearance of Brk binding sites in the *dad* gene completed the network, opening the way to precise decoding of the Dpp gradient over an extended range. Additional feedback has evolved to ensure robustness of BMP signalling in insects and other taxa, and it will be interesting to determine their evolutionary history. We suggest that the Brk circuit is special in that it seemingly operates only in winged insects. Although our work does not inform the debate about the anatomical origin of insect wings (*2,69,3*), it suggests that the Brk module may have been instrumental in their development.

## Acknowledgments

We are grateful to Richard Durbin and Shane McCarthy from the Darwin Tree of Life project for generously sharing the whole-genome sequence of *Thermobia domestica*. We also wish to express our thanks to Siegfried Roth and Matthias Pechmann for their invaluable advice on breeding and experimental techniques with *Thermobia domestica*, and to Aurelio Teleman for kindly providing us with the anti-Brk antibody. We also thank James DiFrisco for discussions. The Advanced Light Microscopy Facility (CALM) at the Francis Crick Institute provided assistance with imaging, and Joachim Kurth from the Crick’s Fly Facility performed the *Drosophila* injections.

The initial phase of this work was funded by a Wellcome Trust Investigator Award to JPV (206341/Z/17/Z). The project was completed with a collaborative grant from the Wellcome trust to JPV and Guillaume Salbreux (223133/Z/21/Z) and core funding from the Francis Crick Institute to JPV (FC001204). The Francis Crick Institute receives its core funding from Cancer Research UK, the UK Medical Research Council, and the Wellcome Trust. The Alexander von Humboldt Foundation supported Luca Cocconi’s contributions to this project.

## Author Contributions

A.H., L.C., and J.-P.V. conceived the study and planned the experiments. A.H. and C.A. carried out the experiments.

A.H. and B. N.-M. analysed the data. L.C., B.N.-M. and G.S. developed the theoretical formalism. L.C. and B.N.-M. performed the numerical simulations. J.-P.V., A.H., L.C. and G.S. wrote the paper with input from all authors.

## Competing Interests

The authors declare that they have no competing interests.

## Supplementary Materials

Materials and Methods

Supplementary Text

Figs. S1 to S10

Table S5

References *(70-81)*

Tables S1-S4

Data S1

## Supplementary Materials For Insect wings arose with a genetic circuit that extents the useful range of a BMP morphogen

## 1 Materials and Methods

### 1.1 Drosophila strains

A list of all genotypes analysed in this study is provided in Table S4. The following stocks were obtained from KYOTO Stock Center or Bloomington Drosophila Stock Center (BDSC): ubi*>*Tkv-HA (DGRC: 118137), dad*>*lacZ or dad^j1E4^ (BDSC, 10305), UAS*>*dad^RNAi^ (BDSC, 33759), brk*>*lacZ or brk^XA^ (BDSC, 58792), tkv*>*lacZ (BDSC, 11191) and ap^MI01996^ (BDSC, 42297). The following fly lines are gifts from various labs: GFP-Dpp (Thomas Kornberg lab, UCSF), UAS*>*tkv (Michael O’Connor lab, University of Minnesota), Tkv-GFP and FRT-Tkv-3xHA-FRT (Markus Affolter lab, University of Basel), dad^4^ *>*EGFPnls and UAS*>*Brk (Giorgos Pyrowolakis lab, University of Freiburg), Omb-GFP and ubi*>*DBD-Mag (Yohannes Bellaïche, Institut Curie), and Sal-GFP (Frank Schnorrer, IBDM). The FRT-HA-Dpp-FRT was previously generated by our lab (*12*).

### 1.2 Transgenes

#### 1.2.1 Generation of various *brk* alleles

The *brk* transcriptional reporter *brk[KO; mCherry]* was generated via CRISPR-Cas9-mediated homology-directed repair. The *brk* gene is intron-less. Two guide RNA-encoding sequences targeting 124 bp upstream of the start codon and 8 bp upstream of the stop codon of the brk coding region were cloned into the pCFD4 vector (Addgene, 49411). Homology arms flanking the *brk* coding region were cloned into the pTV-attP-Pax-mCherry targeting vector. The resulting plasmids were co-injected into a nos-Cas9 strain using in-house fly facilities. The *brk* mutant candidates were identified by *pax-mCherry* expression. Furthermore, since mCherry was sensitive to upstream *brk* regulatory sequences, the strain also served as a *brk* transcriptional reporter.

Subsequently, ФC31-mediated integration into the attP site was used to generate the following lines: brk*>*Gal4, Brk-GFP, FRT-Brk-V5-FRT, Brk^Td^-GFP and FRT-Brk-V5-FRT-Brk^Td^-GFP by cloning the corresponding coding sequences into the pRIV-attB-lox-pax-GFP-lox vector. For the Brk-GFP, Brk-V5 and Brk^Td^-GFP constructs, the coding sequences were flanked by 124 bp of 5’UTR upstream of the start codon and the entire 3’UTR. Integrated candidates were identified by pax-Cherry and pax-GFP double fluorescence. Additionally, pax-GFP served as a *brk* transcriptional reporter in the FRT-Brk-V5-FRT line upon Flp-induced excision.

#### 1.2.2 Generation of UAS*>*Brk^Td^V5

The *Thermobia brk* coding sequence was PCR amplified from genomic DNA (as described below) and subsequently cloned into the pJFRC81-10XUAS vector (Addgene, 36432). The resulting plasmid was injected into a *Drosophila* line carrying the PCary-PattP2 docking site at 68A4 and expressing nos-ФC31 integrase. Successful candidates were identified through the expression of the mini-white marker.

#### 1.2.3 Generation of endogenous GFP-tagged Mad

The *mad[KO; gfp-mad]* allele was generated using a strategy similar to the one described above. Guide RNAs were designed to target the first intron of the Mad-RB isoform, 279 bp upstream of the second exon, and 2bp upstream of the stop codon. The *mad* mutant was created by excising the region between these two target sites. The missing region was then reincorporated, with the addition of EGFP at the beginning of the second exon, in frame with the upstream start codon of the Mad-PB translation isoform.

#### 1.2.4 Generation of the optogenetic Gal4 drivers

The ubi*>*shineGal4sTg is a modified version of the ubi*>*Gal4^Mag^ (*70*). In ubi*>*Gal4^Mag^, the ubi*>*Gal4DBD-Mag and ubi*>*Mag-AD constructs are individually inserted into the genome. In contrast, ubi*>*shineGal4sTg (single transgene) is a single construct containing both ubi*>*Gal4DBD-Mag and ubi*>*Mag-AD in an inverted orientation, separated by a gypsy insulator. This construct was cloned into a pCasper-attB vector and inserted into the attP2 docking site. This single insertion is sufficient to drive light-induced Gal4 activity.

The ap*>*Mag-AD line was generated by cloning the Mag-AD sequence into the pBS-KS-attB2-SA(0) vector (Addgene, 62896) and inserting it into the attP site located in the intronic region of the *ap* gene (ap^MI01996^). The resulting ap*>*Mag-AD fly line was crossed with ubi*>*DBD-Mag to drive light-induced Gal4 activity specifically in the dorsal compartment of the wing disc.

### 1.3 Thermobia Rearing

*Thermobia domestica* (firebrats) were reared in plastic containers with ventilation holes in the lids, stored in a 37 ^°^C incubator under constant dark conditions. To maintain approximately 70% humidity, two uncapped Falcon tubes filled with water were placed in each container. The firebrats were fed commercial goldfish flakes. Additionally, stacks of cosmetic cotton pads were placed in the containers to facilitate embryo collection.

### 1.4 Cloning of the *Thermobia brk* gene

The preliminary genomic assembly of *Thermobia domestica* was generously provided by the Darwin Tree of Life (ToL) Project at the Sanger Institute, with the animal sample supplied by the Siegfried Roth lab at the University of Cologne. The assembly exhibits high genome coverage and contiguity, with an average N50 contig length exceeding 2 Mb. Genomic DNA was prepared from a single firebrat adult using the ChargeSwitch gDNA kit (Invitrogen, CS11203). Since the *brk* gene in Thermobia is intron-less, and consists of 1383 bp, its coding region can be directly amplified from gDNA via PCR. The PCR-amplified fragment was sequenced and validated, showing perfect alignment with the ToL assembly sequence. DNA sequences coding for fusion proteins with V5 or GFP tags fused to the C-terminal end of the Brk^Td^ protein were generated and cloned into the desired vectors using Gibson assembly (New England Biolabs, E5510).

### 1.5 Preparation and sequencing of *Thermobia* cDNA

#### RNA preparation

Approximately 30 *Thermobia* embryos at the limb extension stage were collected in a 0.5 ml Eppendorf tube, mixed with 50 µl of Trizol (Invitrogen, 15596026), and homogenized using a pestle. An additional 450 µl of Trizol and 100 µl of chloroform (Sigma Aldrich, C2432) were then added to the mixture, which was vigorously shaken for 30 seconds. The samples were centrifuged at maximum speed at 4 °C for 15 min. The upper phase was carefully collected and combined with an equal volume of isopropanol. RNA isolation was subsequently carried out using RNeasy Mini Kit (Qiagen, 74104).

#### 3’ Rapid Amplification of cDNA ends (RACE)

The *Drosophila* protein sequences of Dpp, Dad and Tkv were used for tblastn searches against the *Thermobia* genome assembly. This allowed us to identify the scaffold regions where the putative gene homologs were located. Gene-specific primers were designed to target the putative exon regions. To acquire sequence information of the 3’ end of the transcripts, reverse transcription (RT) of the isolated RNA was performed using the SuperScript III kit (Invitrogen, 18080400) with anchored oligo d(T) as the RT primer (CCAGTGAGCAGAGTGAC-GAGGACTCGAGCTCAAGCTTTTTTTTTTTTTTTTTVN). PCR amplification of the 3’end of specific transcripts was conducted using gene-specific forward primers (targeting putative exons) and a reverse primer (GAGGACTCGAGCTCAAGC). The amplified cDNA fragments were then cloned into the pMiniT vector (NEB, E1202) and sent for sequencing.

#### 5’ RACE

To obtain sequence information of the 5’ end of the transcripts, reverse transcription was performed using Template Switching RT Enzyme Mix (NEB, M0466). The anchored oligo d(T) (TTTTTTTTTTTTTTTTTTTTVN) was employed as the RT primer, in conjunction with a Template Switching Oligo (TSO: GCTAATCATTG-CAAGCAGTGGTATCAACGCAGAGTACATrGrGrG). Subsequently, PCR amplification of the 5’end of specific transcripts was carrid out using TSO-specific forward primer (CATTGCAAGCAGTGGTATCAAC) along with a gene-specific reverse primer targeting putative exons. By combining the sequence information from both 5’ RACE and 3’ RACE, we reconstructed the full sequences of the *Thermobia dpp, dad*, and *tkv* transcripts.

### 1.6 Immunostaining and HCR *in situ* hybridization

#### Antibody Staining

*Drosophila* wing discs from the desired larval stages were dissected and fixed in 4% formaldehyde (Thermo scientific, 28906) in PBS for 40 min. The samples were then washed in PBS before overnight incubation in primary antibody at 4 ^°^C. The primary antibodies were diluted in PBS with 0.1% Triton X-100 as follows: rabbit anti-pMad (Cell Signaling, 9516, 1:500), mouse anti-HA (Cell Signaling, 2367,1:200), mouse anti-beta-galactosidase (DSHB, 40-1a, 1:500), mouse anti-Wg (DSHB, 4D4, 1:500), mouse anti-Ptc (DSHB, Apa 1, 1:250), mouse anti-V5 (Invitrogen, 92008, 1:500) and guinea pig anti-Brk (gift from Aurelio Teleman lab, DKFZ, 1:500). The following day, the samples were washed three times with 0.1% Tween-20 in PBS (PBST) before proceeding with secondary antibody staining. Secondary antibodies conjugated with Alexa Fluor (Invitrogen) dyes of the desired excitation/emission wavelengths were used at a 1:500 dilution in PBS with 0.1% Triton X-100, along with Hoechst (Thermo Fisher Scientific, H3570) for nuclear staining. The samples were incubated at room temperature for 2 hours. Subsequently, the samples were washed multiple times in PBST prior to mounting.

#### HCR in situ hybridization

HCR probes were synthesized using two methods. One approach was to submit transcript sequences to Molecular Instruments for probe generation. Alternatively, probes were designed using a script developed by the Ö zpolat Lab at WUSTL (available at https://github.com/rwnull/insitu_probe_generator) and then ordered from Integrated DNA Technologies for oPools Oligo Pools production. Multiplexed HCR immunofluorescence and HCR RNA fluorescence *in situ* hybridization was performed using antibodies, buffers, and hairpins produced by Molecular Instruments. The protocol followed was the generic sample in solution protocol (Rev. 3) available at https://files.molecularinstruments.com/MI-Protocol-2IF-RNAFISH-GenericSolution-Rev3.pdf.

### 1.7 BMP signalling inhibition *in vitro* assay

For chemical inhibition of BMP signalling, LDN-193189 dihydrochloride (TOCRIS, 6053) was dissolved in water to make a 2.5 mM stock solution. *Thermobia* embryos were dissected in Schneider’s Medium (Gibco, 21720024) supplemented with 10% fetal bovine serum (FBS) (Gibco, A3840401). The samples were then incubated either with or without the addition of LDN-193189 at a final concentration of 20 µM in Schneider’s Medium with 10% FBS at 37 ^°^C for 1.5 hours before fixation. Samples were subjected to further analysis, including immunostaining and *in situ* hybridization. To measure the rate of pMad dephosphorylation, fly larvae over-expressing Brk with the dorsal compartment-specific light-induced driver were dissected 6 hours post-illumination in Schneider’s Medium with 10% FBS. The dissected larvae were then incubated with 100 µM LDN-193189 at room temperature for various durations (0, 10, 20, 30, 40, 50, and 60 minutes) before fixation and subsequent immunostaining.

### 1.8 Microscopy

Samples were mounted on glass microscope slides (Epredia, X1XER308B) with No. 1.5 coverslips (VWR, 6310147) using a homemade Mowiol mounting medium (https://cshprotocols.cshlp.org/content/2006/1/pdb.rec10255). Microscopy was performed with a Zeiss LSM880 microscope equipped with a Plan-Apochromat 40x/1.3 NA oil-immersion objective. The pixel size was 346 nm, and the image resolution was 1024 × 1024 pixels. For each sample, a stack of images covering the entire apical-basal axis was acquired with 1 µm intervals between each image.

### 1.9 Image analysis

#### Spatial Profiles of protein concentration in wing discs

Wing discs were co-stained with Hoechst for nuclear segmentation and with anti-Wg and anti-Ptc to determine spatial coordinates (see Fig. S1). Nuclei segmentation was achieved using customized Matlab (RRID:SCR 001622) codes, involving Gaussian filtering, adaptive histogram equalization, and binary thresholding, followed by watershed segmentation. For nuclear-localized signals (e.g., pMad, Brk-GFP, lacZ reporters), the mean nuclear intensity was calculated within a defined diameter around the centroids of segmented nuclei. For cytoplasmic or membrane signals (e.g., *brk* transcriptional reporters, GFP-Dpp), mean intensity within the region marked by the blue shade in Fig. S1 was quantified along the A-P axis. Perturbations were carried out in the dorsal compartment when possible, comparing raw gene expression intensity between dorsal and ventral compartments to minimize systematic errors. When cross-individual comparisons were necessary, samples were immunostained and imaged under identical conditions to ensure comparable raw intensity.

#### Transcript counting in Thermobia limb progenitors

Nuclei segmentation of *Thermobia* embryos was performed using a modified Python (RRID:SCR 008394) code based on the Cellpose algorithm. Spatial coordinates within limb progenitors were defined by setting the distal end of the limb to zero and manually drawing a central midline along the proximal-distal (P-D) axis. Nuclear centroids were projected onto this midline to determine their position in µm along the P-D axis. pMad nuclear intensity was calculated as the mean intensity within the contour of each segmented nucleus. HCR signals were segmented using a modified Python code based on Big-FISH smFISH image processing package (*71*). Each segmented transcript was assigned to the nearest nuclear centroid. The P-D axis was divided into 10 µm bins, and the mean and standard deviation of pMad intensity and transcript count per cell were computed for each bin.

### 1.10 Expected *a posteriori* Distribution

To quantify and visualise uncertainty in implied position we employ the expected *a posteriori* distribution (*26*). To calculate this, the conditional probability density of gene expression levels or signalling activities *g* given position *x* is first approximated by a Gaussian distribution

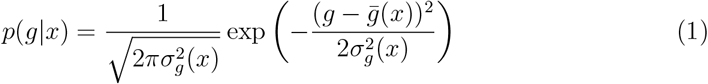

where 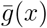 and *σ*_*g*_(*x*) are the empirical mean and standard deviation of the gene expression level or signalling activities *across replicates*, respectively. We note that such an approximation has been previously used in analysis of positional uncertainty in morphogen gradients (*72*). To measure gene expression levels or signalling activities *in each replicate*, nuclei in a thin strip along the D-V boundary were segmented and binned over 50 uniform bins in the range *x/L* ∈ [0, 1], with gene expression levels or signalling activities given by the empirical mean value of the *M* nuclei in a given bin (*M* ≈ 8 in all cases). These average values inside each bin were then used to evaluate *σ*_*g*_ across replicates.

We now briefly discuss the biological interpretation of this measured variability. For any given bin *b* = 1, …, 50 and denoting 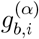 the gene expression level or signalling activity of the *i*-th cell (*i* = 1, …, *M*) of the *α*-th sample, we consider the following covariance, ignoring cross-correlations for simplicity:

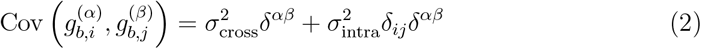

with *δ* the Kroeneker delta, where 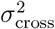 and 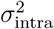 denote contributions to the single-cell signal variance originating from cross-sample and intra-sample variability, respectively.

We intuitively expect *σ*_cross_ to be associated to pattern robustness across individuals, and *σ*_intra_ to pattern precision within one individual. The intra-sample variability however may stem from a combination of “relevant” biological variation, but also from “irrelevant” measurement error and systematic variability due to cell-cycle dynamics, the relative magnitude of which is challenging to assess. Based on Eq. (2), we have that the empirical cross-sample variance can be written as 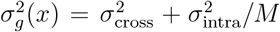.

Thus, *σ*_*g*_(*x*) is an upper bound on the cross sample variability and a lower bound on the total single-cell variability that would be obtained by considering variability among individual nuclei, including measurement error. Given the similar conditions and measurement techniques used in experiments with internal controls, the measurement error is likely to be of comparable magnitude and structure, allowing for relative comparisons to still be valid.

The conditional distribution (1) of gene expression levels or signalling activities is then employed to find the posterior distribution of the implied position *x*^*^ through Bayes’ theorem:

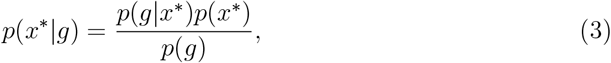

where *p*(*g*) is the probability distribution over gene expression levels or signalling activities and *p*(*x*^*^) is the prior distribution, which we take to be uniform over [0, *L*]. The expected *a posteriori* distribution is defined as

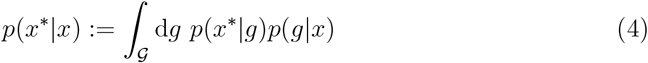

and captures the average likelihood that could be assigned by a cell at true position *x* to an implied position *x*^*^ upon observation of its own gene expression level or signalling activity, if provided with knowledge of the joint distribution *p*(*g, x*). We remark that the definition (4) is agnostic to the specific choice of decoder, i.e. to the rule that a cell might subsequently follow when “committing” to a specific guess for *x*^*^.

Eqs. (1) and (4) can be extended to a vector of multiple gene expression levels or signalling activities 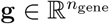 using a multivariate Gaussian approximation

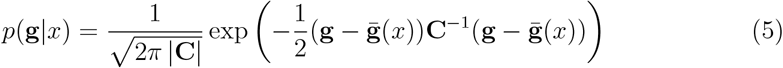

where **C**_*ij*_ Ξ Cov(*g*_i_(*x*), *g*_j_(*x*)) denotes the covariance matrix, whence

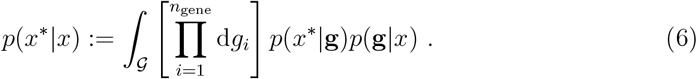

Note that the expected *a posteriori* distribution as defined in Eq. (4) is closely related to the decoding map 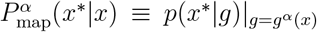 introduced by Petkova et al. (*27*), where *α* indexed replicates. In particular, assuming that the Gaussian approximation is accurate, the expected *a posteriori* distribution is the limit, as the number *N* of replicates goes to infinity, of the mean decoding map:

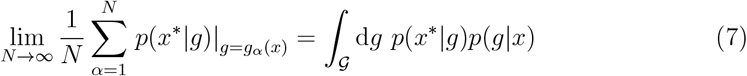

since *g*^α^ is distributed according to *p*(*g*|*x*).

### 1.11 Positional Information Density

We define the *Gene Information Gain* as

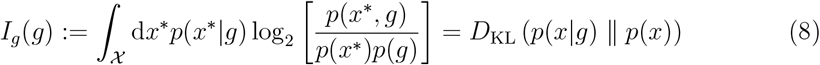

where *D*_KL_ (*P* ‖ *Q*) denotes the Kullback-Leibler divergence between the distributions *P* and *Q* (*73*). This quantity, measured in bits, represents the reduction in uncertainty about the true position *x* after observing a specific value *g* of gene expression level or signalling activity. The mutual information between position and gene expression/signalling activity, often referred to as positional information in this context (*72*), is given by the expected value of the information gain (8) over *g*,

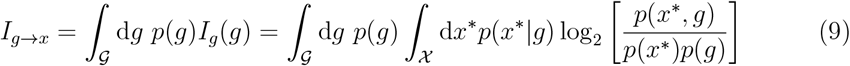

We further define the *Positional Information Density* (PID) as a conditional analog of the mutual information (9),

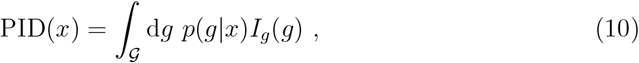

which can be straightforwardly generalised to a set of gene expression levels or signalling activities **g**, along the lines of Eq. (6). The PID quantifies the average reduction in uncertainty about the position *x* given observations of gene expression level or signalling activity *g* specifically at that position. We refer to it as the “positional information density” because it is a local measure of information content, and its integral over all *x*, weighted by the prior distribution *p*(*x*), recovers the total positional information. We use this quantity to evaluate positional information returned independently by Dpp signalling measured by pMad or *brk* expression level (Fig. 2E, F) or by their combined effect (Fig. 2K, L).

While the PID is an information-theoretic measure and does not directly translate into a spatial scale (such as a “positional error”), it is helpful to consider its intepretation in a simplified case. Suppose the posterior distribution *p*(*x*|*g*) = 𝟙 _Ω(*g*)_(*x*) is a normalised indicator function over one of 2^*N*^ equally sized and distinct (though not necessarily continuous) subsets 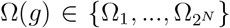 of the interval [0, *L*], where |Ω (*g*)| = *L/*2 for any *g*. In this scenario, we have *I*_*g*_(*g*) = *N* bits, and therefore PID(*x*) = *N* bits as well for any *p*(*g*|*x*). Thus, for this particular case, a positional information density of *N* bits allows cells at position *x* to uniquely localise themselves within one of 2^*N*^ equally sized subdomains of the interval [0, *L*]. For intuition, one may associate a length scale 𝓁_PID_ with the PID, defined as 𝓁_PID_(*x*) Ξ *L/*2^PID(x)^. Note that, unlike the positional error of Ref. (*27*), this heuristic length scale is decoder-agnostic.

### 1.12 Identification of Brk protein homologs

To identify *brk* genes in various genomes, the protein sequence of *Drosophila melanogaster* (*Dmel*) Brk (accession: NP 511069; 704 amino acids) was used as a query in TBLASTN searches on the NCBI database. High-coverage genomic sequences or assemblies with large N50 contig sizes were selected, covering the orders Diptera, Lepidoptera, Coleoptera, Hymenoptera, Orthoptera, Ephemeroptera, Odonata, and Zygentoma. All searches yielded at least one BLAST hit, typically within single proteincoding regions from start to stop codons, indicating that the *brk* gene in most species is intron-less, with the exception of a few species in Coleoptera and Orthoptera (see Fig. S4B). This was confirmed by available transcriptomic data. Protein sequences from these single-exon coding regions were then aligned using MUSCLE v3.8.1551 with default parameters. The DNA-binding domain of *Dmel* Brk showed significant alignment across all identified Brk homologs. Co-repressor recruitment motifs of Brk were manually searched and annotated by identifying the amino acid sequences PMDLSLG for the CtBP interaction motif and FKPY for the Groucho interaction motif (*39*).

For non-insect hexapods where high-quality genome assemblies were not available in the NCBI database, TBLASTN searches were conducted in the Ensembl Metazoa and Ensembl Rapid Release databases using *Dmel* Brk as the query within the taxonomic group “Entomobryomorpha.” Transcripts containing hits were analyzed as described above.

To investigate the presence of the *brk* gene in hexapod outgroups, whole-genome proteome data from various taxonomic groups (one species each from Arachnida, Branchiopoda, Chilopoda, Copepoda, Malacostraca, Merostomata, Thecostraca, and seven hexapod species, including *Dmel*) were downloaded from the Ensembl Metazoa database. These data were analysed using OrthoFinder v2.5.4 to generate orthogroups. The orthogroup containing *Dmel* Brk included Brk orthologs from seven hexapod species but none from outgroups, confirming that *brk* is a hexapod-specific gene.

### 1.13 Phylogenetic analysis of Brk protein homologs

To construct the phylogenetic tree of Brk homologs, Brk DNA-binding domains (DBDs) from hexapod species were identified through significant sequence alignment to the *Dmel* Brk DBD. These DBD sequences were aligned using the MUSCLE algorithm (*74*). The resulting protein alignment was used to construct a neighbor-joining (NJ) tree based on BLOSUM62 substitution matrices. The NJ tree was generated using Jalview software (*75*), providing a visual representation of the evolutionary relationships among the Brk DBDs.

### 1.14 Identification of putative silencer regulatory elements upstream of *brk* homologs

To identify putative silencer motifs in *brk* gene homologs, we analyzed a 20 kb genomic region upstream of the transcription start site for each homolog. If the transcript was not annotated, the region was taken as 20 kb upstream of the translation start site of the Brk protein homologs. This region was searched for the consensus DNA sequence GRCGNCNNNNNGTCT (where N represents any nucleotide, and R represents A or G), known to be recognised by the Mad-Medea-Schnurri complex, mediating transcriptional repression (*68, 76*). All identified putative silencer motifs were recorded in Table S2. Representative configurations of the putative silencer motifs across species within each order were illustrated in Fig. S5I.

### 1.15 Identification of putative Brk-binding sites in *dad* homologs

To identify putative Brk-binding sites in *dad* homologs, we searched for the DNA consensus sequence GGCGCCANNNNGNCV (where V represents A, C, or G), which has been shown to be competitively bound by the pMad-Medea activation complex or Brk in the long intronic regions of *dad* homologs (*31*). Identified putative Brk-binding sites that strictly matched the consensus sequence, as well as those with relaxed linker sequences (GGCGCCNNNNNGNCV) or relaxed Mad binding sequences (GGCGY-CANNNNGNCV), were recorded in Table S3. Schematics of the putative Brk-binding sites, with one representative species from each order, were illustrated in Fig. S5J.

## 2 Supplementary Text

In this supplementary text, we present an experimentally-informed model describing the establishment of a steady-state Dpp/BMP morphogen gradient along the A-P axis of the *Drosophila* wing disc. Our model accounts for cell-autonomous signalling dynamics, via the inclusion of the repressive target of pMad, Brk, which in turn modulates Mad phosphorylation by acting as a transcriptional repressor of the Mad competitor, Dad. This incoherent feedback motif plays a key role in extending the Dpp morphogen gradient, improving its readability in distal regions. The model additionally accounts for non-autonomous effects, via indirect transcriptional upregulation of the receptor Tkv by Brk, leading to enhanced signalling (and ligand degradation through receptor-mediated internalisation) in the medial region of the tissue. As we will see, while highly non-trivial, this second incoherent feedback element appears to have relatively negligible effects on the steady-state morphogen gradient. The parameters appearing in the definition of the model are fitted to a number of experiments dissecting the gene regulatory network downstream of Dpp signalling, as detailed in the following. Having fixed these parameters, we observe good quantitative agreement between model and experimental gradients under wild-type as well as mutant conditions.

### 2.1 Model definition

Our model of Dpp signalling consists of two modules. The first module describes extracellular Dpp transport along the A-P axis via a one-dimensional diffusion-degradation equation of the form,

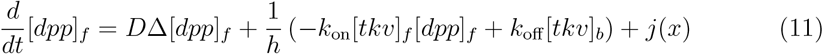

where [*dpp*]_*f*_ denotes the concentration of free extracellular Dpp, in units of inverse volume, *D* is the effective diffusivity, *h* is the width of the intercellular space and *k*_on_ and *k*_off_ are the binding and unbinding rates to/from the receptor Tkv. The membrane concentrations of free and bound receptors are denoted [*tkv*]_*f*_ and [*tkv*]_*b*_, respectively. Finally, *j* is a space-dependent source term vanishing everywhere outside the dpp source stripe. Here, we ignore spontaneous Dpp degradation and leakage (*77*) for the sake of simplicity.

The second module describes the intracellular signalling dynamics driven by Dpp binding to Tkv, which promotes phosphorylation of Mad when the intracellular domain of Tkv is not occupied by the Mad competitor Dad,

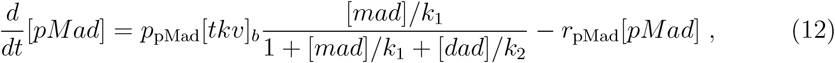

where *p*_pMad_ is the phosphorylation rate per unit of bound receptor membrane density and *r*_pMad_ is the dephosphorylation rate. The fraction appearing in the first term in the right hand side of Eq. (12) captures the probability that the intracellular binding site of Tkv is occupied by Mad, given dissociation constants *k*_1_ and *k*_2_ for Tkv of Mad and Dad, respectively.

Phosphorylated Mad (pMad) acts as a transcription factor (TF), repressing *brk* and activating *dad*. Meanwhile, Brk itself acts as a repressor, transcriptionally downregulating *dad* and most likely upregulating *tkv* in an indirect manner. Moreover, it cooperates with pMad to repress its own transcription (although, as discussed below, we neglect this effect here). These regulatory interactions are captured by the following set of equations,

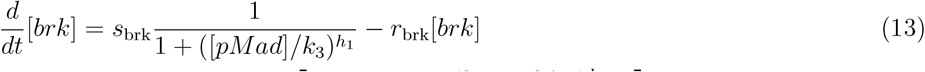

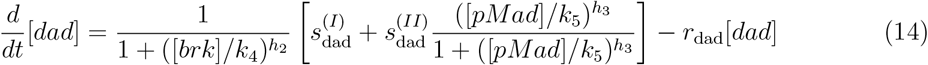

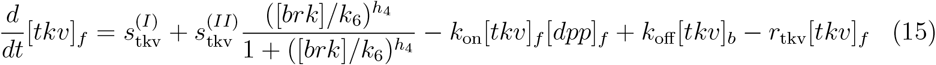

and schematised in Fig. 3H. Finally, for the dynamics of the concentration of bound receptors we write

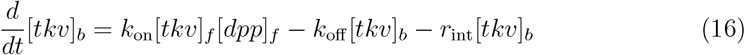

where in general *r*_int_ ≠ *r*_tkv_ to account for the possibility of active internalisation of the signalling complex. In the above equations, [•] once again denotes concentrations, in units of inverse volume or surface area, of the relevant species. Positive and negative interactions are modelled by a-priori nonlinear Hill functions with exponent *h*_•_ and characteristic concentration *k*_•_. Parameters related to protein production, denoted *s*_•_, can appear with different superscripts within the same equation to distinguish baseline vs activated production. Previous literature has argued for a direct transcriptional downregulation of *tkv* by Dpp signalling, for which we have found no empirical evidence and which we have thus ignored.

Note that we have not written an explicit dynamical equation for the concentration of unphosphorylated Mad, [*mad*]: since total Mad, [*mad*_tot_] = [*mad*] + [*pMad*], is assumed constant in time and space and assuming for simplicity that [*pMad*]*/*[*mad*_tot_] ≪ 1, we can replace [*mad*] by the constant concentration [*mad*_tot_] in the expression for the phosphorylation flux. This assumption is generally valid in the distal region of the gradient, where Mad phosphorylation is low, which is the region of interest in the present study.

#### 2.1.1 Steady-state equations

At steady-state, *d*[•]*/dt* = 0 for all species. Eq. (16) then reduces to

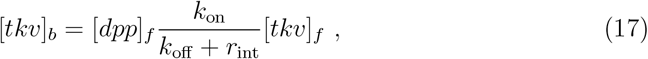

allowing us to eliminate [*tkv*]_*b*_ in favour of [*tkv*]_*f*_ in all other equations. Substituting into the steady-state form of the diffusion equation (11), we obtain

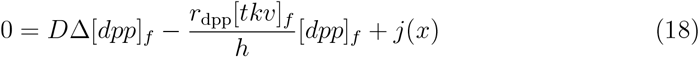

where *r*_dpp_ = *r*_int_*k*_on_*/*(*k*_oα_ + *r*_int_), which now features an effective inhomogeneous degradation term proportional to the local concentration of free receptors. Analogous substitution into Eq. (15) for the concentration of free receptors yields

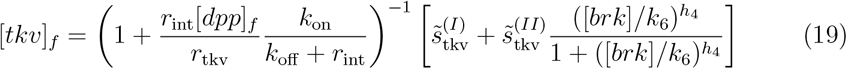

where 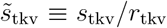. Assuming that we are working far from saturation, specifically *r*_int_[*tkb*]_*b*_*/*(*r*_tkv_[*tkv*]_*f*_) ≪1, the dependence on [*dpp*]_*f*_ in the prefactor of Eq. (19) can be neglected and we are left with the simpler expression

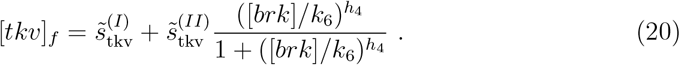

This approximation is always valid sufficiently far from the Dpp source. A conservative estimate of 40*µm* for the distance beyond which it applies can be obtained by noting that at steady-state [*tkv*]_*b*_*/*[*tkv*]_*f*_ ∝ [*dpp*]_*f*_ and drawing on the characteristic Dpp gradient decay length *𝓁* ≃ 10*µm*, discussed below in Sec. 2.3, together with *r*_int_*/r*_tkv_ ≃ 10^2^ (*77*) and assuming 50% receptor occupancy at the A-P boundary.

Finally, we write for the remaining species

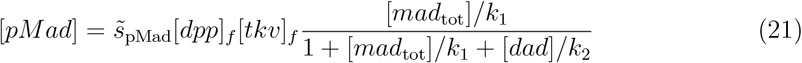

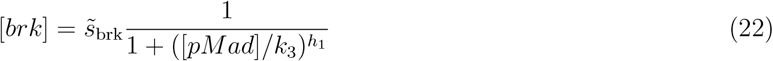

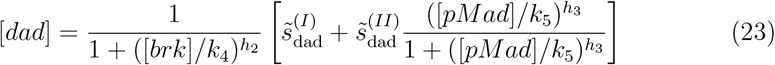

where again 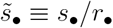, after having defined *s*_pMad_ = *p*_pMad_*k*_on_*/*(*k*_oα_ +*r*_int_). We can solve these nonlinear equations numerically as a function of [*dpp*]_f_ to obtain a set of input-output response curves characterising the cell-autonomous response to a chemostatted external signal, as shown in Fig. 3I-L of the main text. To do so, however, we first need to fix a number of model parameters.

We henceforth drop the subscripts for [*dpp*]_*f*_ and [*tkv*]_*f*_ for the sake of brevity, with the understanding that [*tkv*]_*b*_ will not play an explicit role in the following.

#### 2.1.2 The *brk* mutant

Due to the importance of Brk as a factor regulating the transcription of *tkv* and *dad*, a genetic perturbation that played an essential role in dissecting the associated gene regulatory network is the genetic ablation of the DNA region coding for the Brk protein itself. We refer to this genotype as a *brk* mutant. At a mathematical level, it corresponds to setting *s*_brk_ = 0 in Eq. (13), so that the steady-state equations reduce to

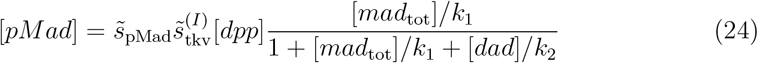

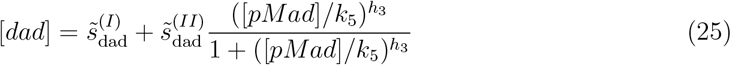

with 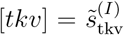 and [*brk*] = 0. Notwithstanding the absence of the Brk protein, it is still possible to measure the rate at which its RNA would have been transcribed using a transcriptional reporter, *brk[KO; mCherry]*, as described in the main text. We can write the steady-state expression of this transcriptional reporter as

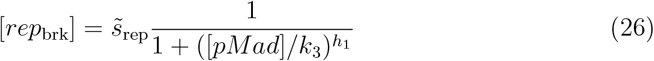

where in general 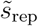 differs from 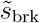 due to the different lifetimes of the two proteins and their respective RNAs.

### 2.2 Parameter fit from cell-autonomous response

The intracellular module of our mathematical model, as formalised by Eqs. (12)-(16), involves a relatively large number of parameters. Fortunately, as far as the steady-states (21)-(20) are concerned, not all are independent, *e*.*g*. doubling all production, *s*_•_, and degradation, *r*_•_, rates simultaneously leaves the steady-state solutions unchanged. Furthermore, we can work in arbitrary units of concentration such that a subset of parameters can be set to unity without loss of generality. For example, since [*brk*] only ever appears as a ratio involving *k*_4,6_, we can set 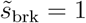 with the understanding that fitted values *k*_4,6_ are now in units of 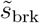. The parameters that cannot be removed in this way are fitted to experimental data, as described below.

#### 2.2.1 Effective input-output response curve for pMad-*brk* branch

From images of wild-type (WT) and *brk* mutant wing discs, it was possible to extract input-output response curves relating intracellular pMad concentration and *brk* transcription, inferred from the transcriptional reporter, as shown in Fig. S7A. Crucially, the possibility of inducing *brk* mutation only in the dorsal compartment meant that the two response curves could be compared directly. The result for the *brk* mutant background is a curve that can be fitted to Eq. (26) to extract *h*_1_ = 1.60 ± 0.04 and *k*_3_ = 0.0459 ± 8 × 10^−4^, the second being in units of the pMad concentration at the A-P boundary in the WT compartment. The response curve for the WT background displays a slightly sharper decay of *brk* transcription with increasing pMad, suggesting weak *brk* self-repression. Matching of the two curves at vanishing pMad concentration indicates that such self-repression is not direct (consistent with the absence of functional Brk binding sites in the upstream *brk* regulatory sequence) but might instead originate from a stabilisation of the pMad-Medea complex at the promoter (*78*). Since the specific molecular mechanism underlying Brk self-repression is at present unknown and considering its comparatively weak impact, we decided to ignore it in our model, cf. Eq. (13).

#### 2.2.2 Effective input-output response curve for pMad-*dad* branch

A similar procedure can be followed to fit the model parameters appearing in Eq. (23) for the steady-state Dad concentration to the input-output response curve relating pMad and Dad, obtained by combining the data shown in Figs. 2C and 3D. To do this, we first use the pMad-*brk* response relation determined in the previous section to eliminate [*brk*] and express the right hand side of Eq. (23) purely as a function of [*pMad*].

The empirical response curve is shown in Fig. S7B. We thus obtain *h*_2_ = 3.0 ± 0.3 and *h*_3_ = 3.1 ± 0.4 for the Hill exponents, as well as 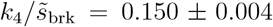 and *k*_5_ = 0.60 ± 0.02, again in units of the pMad concentration at the A-P boundary in the WT compartment. Comparing *dad* expression in the proximal region ([*pMad*] ≃ *k*_5_) vs the distal end of the gradient ([*pMad*] ≃ 0) in the *brk* mutant compartment (Fig. 3D) we can also estimate 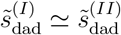. Similarly, by comparing the median vs distal signals from simultaneous measurements of *dad* and *brk* transcriptional reporters, Fig. S8A, and assuming comparable degradation rates for the two proteins, we can estimate 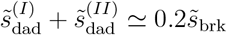.

#### 2.2.3 Effective input-output response curve for pMad-*tkv* branch

Finally, the parameters appearing in Eq. (20) can be fitted by considering the input-output response relating intracellular pMad and extracellular Tkv concentrations, the latter obtained by immunofluorescence. To do so, we first use Eq. (22) (with the fitted parameters obtained in Sec. 2.2.1) to replace [*brk*] in Eq. (20) with a function of [*pMad*] only. It should be noted that, in reality, the expression of [*tkv*] is not homogeneous even in the absence of Brk protein (see Fig. 3G), when the second term in the right-hand side of Eq. (20) vanishes. This is due to inhomogeneous Hedgehog (Hh) signalling upstream of *tkv* transcription, which we ignore here for the sake of simplicity and on the basis of its relatively minor effect. To circumvent this complication, we fit Eq. (20) in the wild-type case by first subtracting the inhomogeneous *tkv* baseline in the *brk* mutant background, see Fig. S7C. We thus obtain *h*_4_ = 5 ± 1 and 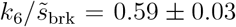. By comparing *tkv* expression in the distal region of the gradient across wild-type and *brk* mutant we can also estimate 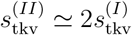.

#### 2.2.4 Competitive inhibition of Mad phosphorylation by Dad

The inhibition of Mad phosophorylation by Dad is modelled in Eq. (21) in terms of competitive binding for the same binding site within the intracellular domain of Tkv. We assume that depletion of free Dad and Mad by binding to Tkv is negligible. Thanks to our other assumption that the phosphorylated fraction of total Mad is negligible, the concentration and dissociation constant of Mad for said binding site only ever appear in combination, so that [*mad*_tot_]*/k*_1_ constitutes a single free parameter. Nevertheless, we can independently estimate 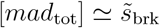 by noticing that fluorescence levels of Brk-GFP and GFP-Mad are comparable in the distal region of the gradient, Fig. S8B. As far as the relative values of the characteristic concentrations *k*_1_ and *k*_2_ are concerned, these cannot be estimated directly from the available data. However, existing qualitative analysis (*58*) (and indeed our own results presented below) supports the assumption that *k*_2_ ≪ *k*_1_. If we further assume that the probability of Tkv not being occupied by either Dad or Mad is negligible, *i*.*e*. demanding 1 + [*mad*_tot_]*/k*_1_ + [*dad*]*/k*_2_ ≃ [*mad*_tot_]*/k*_1_ + [*dad*]*/k*_2_, we are left with a single free parameter proportional to the ratio *k*_2_*/k*_1_. The validity of this last assumption was checked *a posteriori* upon comparison with empirical gradients.

### 2.3 Establishing gradients in space

The relaxation of the extracellular Dpp gradient to its steady-state shape is determined by the diffusion-degradation dynamics in Eq. (11), which depends at every point *x* ∈ [0, *L*] on the local membrane concentration of Tkv. Here, *x* = 0 corresponds to the A-P boundary, while *x* = *L* with *L* ≃ 100*µm* denotes the impermeable boundary at the posterior end of the tissue, where we impose a no flux boundary condition. In turn, [*tkv*] depends on Dpp signalling via the gene regulatory network described above. In this sense, and in contrast with the traditional picture of morphogen gradients simply propagating positional information from a source region into a homogeneous tissue, tissue-level positional information in the wing disc is just as much an emergent property of morphogen-mediated interactions between cells.

In keeping with the parameter fit discussed in the previous section, we work in units such that, under wild-type conditions, [*pMad*] = 1 at *x* = 0, which provides us with the the second boundary condition. When considering other genotypes, unless otherwise specified we instead impose a constant flux boundary at *x* = 0 such that the Dpp flux into the posterior compartment matches that of the wild-type disc, which we extract from the solution of the wild-type gradient.

Rather than solving Eqs. (11)-(16) simultaneously, we instead solve Eq. (11) using the instantaneous steady-state values (21)-(20). This is mathematically equivalent to assuming a separation of time scales between fast intracellular and slow extracellular dynamics, although the validity of this assumption does not affect the steady-state properties of the gradient. *D* and *r*_dpp_*/h* are now left as the only parameters proportional to a rate. We thus rescale time such that *D* = 1 and set 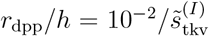, which is consistent with the experimental observation that the characteristic length (*79*) of the (exponential) dpp gradient estimated in the proximal region is

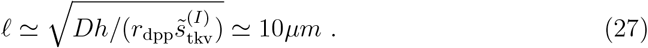

Having fixed all parameters of the model except for the ratio *k*_2_*/k*_1_, we can now solve numerically for the steady-state concentrations profiles of all species of interest. We do this for both wild-type and *brk* mutant conditions, the latter’s governing equations being given in Sec. 2.1.2. Note that up to this point we never fit directly to the spatial gradients, instead extracting each free parameter from cell-autonomous, input-output response curves, as described in the previous section. Nevertheless, and notwithstanding the various approximations made along the way, we find a remarkable quantitative agreement between the gradients predicted by our model and those observed experimentally as long as *k*_2_*/k*_1_ ⪅10^—2^, see Fig. S7D, consistently with previous evidence that *k*_2_ ≪ *k*_1_.

For the sake of clarity, we summarise the fitted parameters (or parameter combinations) mentioned in the previous sections in Table S5. These were used to generate the model curves in the relevant figures, unless specified otherwise. The similarity between model and experimental results supports our conclusion that the components of the gene regulatory network downstream of dpp that were identified in the present work are *su cient* to explain both the extension of the wild-type pMad gradient and the smoothing of the brk inverse gradient, as compared to the respective brk mutant counterparts. In other words, while interactions with components not currently accounted for cannot be excluded in principle, they are not necessary to understand these effects.

### 2.4 Genetic perturbations, *in silico* and *in vivo*

Having established the validity of the model in wild-type and *brk* mutant conditions, we now use it to ask questions about the relative importance of the two key incoherent feedback motifs appearing in the Dpp signalling network, namely *dad* downregulation and *tkv* upregulation by Brk. We do this by artificially ‘switching off’ either of the two regulatory interactions and observing the impact of this ‘mutation’ on the steadystate gradients. We then further verify the robustness of the model by comparing its predictions with experimental data under two rescue genotypes with *brk* mutant background, where the Brk protein is replaced either by DNA coding for Tkv (thus effectively reintroducing the positive interaction by which Brk activates *tkv*) or for *dad*^*RNAi*^ (effectively reintroducing the negative interaction by which Brk represses *dad*).

#### 2.4.1 Relative importance of the incoherent feedback motifs

In order to determine the relative importance of the two incoherent feedback motifs (Brk repressing *dad* and Brk activating *tkv*) we modify our model to remove each in turn. Upon deactivating the repression of *dad* by Brk while maintaining Brk’s activating effect on *tkv*, the steady-state equations for [*dad*] and [*tkv*] become

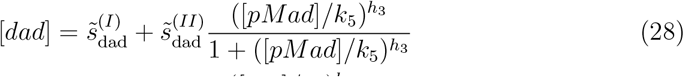

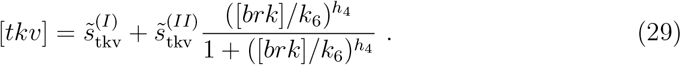

In this case the model produces pMad and Brk gradients that match those of the *brk* mutant genotype, see Fig. S7E. In particular, both pMad gradient extension and Brk inverse gradient smoothing are abrogated in this case. Vice versa, deactivating the positive effect of Brk on *tkv* while maintaining its repressive effect on *dad* corresponds to modifying the steady-state equations as

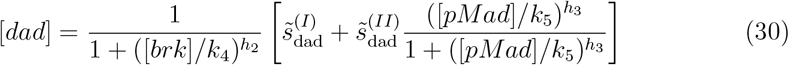

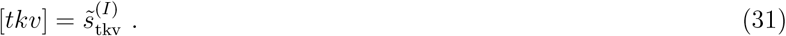

The resulting steady-state gradients of both pMad and Brk closely resemble those of the wild-type genotype, although the *tkv* expression profile is now homogeneous (See Fig. S7F). This observation supports the conclusion that *dad* repression by Brk constitutes the dominant mechanism in establishing robust, spatially extended gradients as required for the patterning of the wing disc.

##### 2.4.1.1 *ubi>tkv* genotype

We approached the condition modelled by Eqs. (30) and (31), albeit imperfectly, by genetically ablating the endogenous *tkv* and providing Tkv ubiquitously from a ubiquitin-driven transgene (*ubi>tkv*). This genotype, denoted *ubi>tkv*, diverges from the ideal one described above in two ways: firstly, the uniformly expressed *tkv* level might differ from that in a wild-type disc where *tkv* upregulation by Brk is abolished (*i,e*.,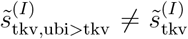); secondly, the ubiquitin driver remains active within the Dpp source stripe, unlike the endogenous *tkv* which is transcriptionally repressed by Hh signalling. This enhances Dpp uptake in the source, thus reducing the net flux of Dpp into the posterior compartment, which constitutes a boundary condition in our problem, as discussed in Sec. 2.3. Moreover, pMad signalling has been shown to negatively regulate Dpp production, further reducing the morphogen outflux from the source stripe.

To quantify the magnitude of the first effect, we compared the relative transcription driven by a *ubiquitin* promoter with that from the endogenous *tkv* promoter by co-expressing an endogenously HA-tagged conditional *tkv* allele and a *ubi>tkv-HA* transgene in the same wing disc. We specifically expressed Flp in the dorsal compartment to eliminate endogenous Tkv-HA there, allowing us to compare HA immunofluorescence in the dorsal (*ubi>tkv-HA* alone) versus the ventral (*ubi>tkv-HA* + one copy endogenous *tkv-HA*) compartments. We observe a twofold ratio between the two at *x/L* = 0.7 (see Fig. S9A-B), likely a slight underestimate due to the impossibility of systematically subtracting the autofluorescence background. Thus, 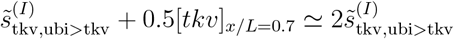, the factor of 0.5 coming from only one copy of endogenous *tkv* being tagged. Combining this result with the fact that, at the same location, Tkv expression in a *brk* mutant background is approximately one-third of that in wild type (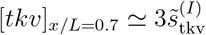, see Fig. 3G), we conclude that Tkv expression levels in *ubi>tkv* are largely comparable to the baseline expression in wild type, whence we set 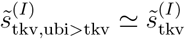.

The reduction in the Dpp diffusive flux across the A-P boundary was estimated by comparing the spatial distribution of endogenous GFP-Dpp in wild-type discs to that in discs expressing Tkv from the *ubi>tkv* transgene in addition to endogenous Tkv. Assuming that the fluorescence from GFP-Dpp is quickly quenched in endocytic vesicles and that the fraction of extracellular GFP-Dpp is small, the integral over the source area of the fluorescence signal provides a measure of the difference in Dpp production. The two assumptions, amounting to the statement that the contribution to the total fluorescence signal originating from extracellular and endocytosed GFP-Dpp is negligible compared to that of pre-secretory GFP-Dpp, are supported by the fact that GFP-Dpp signal is much lower in receiving cells than in producing cells. Accordingly, we estimate an 11% reduction in total Dpp production (Fig. S9C-D). From this and the knowledge that the degradation rate increases due to the increased *tkv* expression in the source region in *ubi>tkv* compared to WT, we can obtain an upper bound to the change in Dpp diffusive flux at the A-P axis relative to the wild type, specifically *j*_ubi_*/j*_WT_. 0.89. The true value should approach the bound when the change in Dpp degradation in the source across the two genotypes is negligible. Inputting 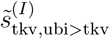 into our modified model without *tkv* activation by Brk, Eqs. (30) and (31), and sweeping over *j*_ubi_*/j*_WT_ from 0.3 to 0.9, we see the best match to experimental profiles is obtained for *j*_ubi_*/j*_WT_ ≃ 0.3. This suggests that enhanced Dpp degradation in the source indeed has a non-negligible effect. In summary, the minimal impact on the establishment of the reverse Brk gradient when replacing endogenous Tkv with uniformly expressed Tkv suggests that the Brk-*tkv* feedback branch is not crucial to this process.

#### 2.4.2 Qualitative comparison to single feedback branch rescue genotypes

Selective deactivation of only one of the two feedback branches is challenging from an experimental point of view. As an alternative approach to exploring their role in isolation, we have constructed two ‘rescue’ genotypes in a *brk* mutant background.

##### 2.4.2.1 *brk>tkv* rescue genotype

In the first genotype, the *brk* promoter drives graded Tkv expression in the *brk* domain using the GAL4/UAS system. We model this by using Eqs. (28) and (29) for the [*dad*] and [*tkv*] dynamics, replacing [*brk*] everywhere by [*GAL*4], which satisfies

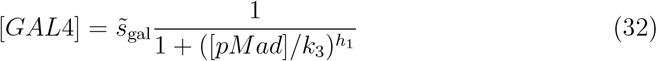

in the spirit of Eq. (26) for the *brk* transcriptional reporter. We further increase *s*^(II)^ by a factor 10 compared to the wild-type value to approximately account for strong expression of GAL4/UAS and set 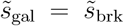 for the sake of simplicity. Since the positive transcriptional input of GAL4 on *tkv* will in general differ in strength from that of endogenous Brk, we evaluate the modified 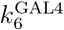 and 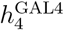 (cf. Eq. (29)) in this genotype by fitting the predicted pMad profile onto the experimental one, shown in Fig. 4C. We thus obtain 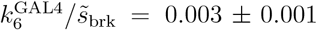 and 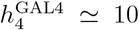, with a large uncertainty on the latter due to poor convergence of the optimizer. Nevertheless implementing these changes in the model leads to an increase in pMad level relative to the *brk* mutant parameter regime and a small increase in the sharpness of the *brk* expression profile, qualitatively matching the experimental results (see Fig. 4C, C’). The failure of *tkv* overexpression in both simulation and experiment to rescue the *brk* profile suggests that *tkv* feedback is insufficient to extend the gradient.

##### 2.4.2.2 *brk>dad^RNAi^* rescue genotype

In the second test genotype, the *brk* promoter drives *dad*^*RNAi*^, inhibiting *dad* expression far from the Dpp source. We model this by using Eqs. (30) and (31) for [*dad*] and [*tkv*], replacing [*brk*] everywhere by [*dadRNAi*], which satisfies

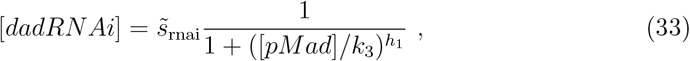

again in a similar vein as Eq. (26). Here, we set 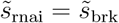 for the sake of simplicity. As previously done for the *bkr>tkv* rescue genotype, we evaluate the modified parameters 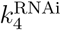 and 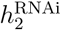 (cf. Eq. (30)) by fitting the predicted pMad profile onto the experimental one, shown in Fig. 4B. In this case, we obtain 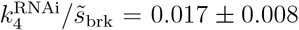 and 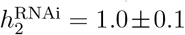. Note that while 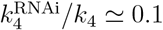 suggests enhanced inhibition (as may be expected), this interpretation relies on the validity of the assumption 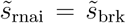 made above. In this case, our model predicts some degree of smoothing of the *brk* inverse gradient (partial rescue), together with amplification of the pMad signal in the medial region, in qualitative agreement with experimental results (Fig. 4B). Once again, these findings support our conclusion that negative auto-regulation mediated by Dad plays a predominant role in the establishment of a shallow *brk* gradient.

#### 2.4.3 *dad* mutant genotype

The experimental evidence presented up to this point supports our conclusion that competitive inhibition of Mad phosphorylation by Dad plays a key role in the establishment of both the pMad and Brk morphogen gradients by differentially sensitising cells to the extracellular signal in the medial region of the gradient. Indeed, both the existing literature (*58*) and our own analysis indicates that Dad might have a much larger affinity than Mad for the intracellular domain of Tkv, and thus exerts a strong inhibitory effect in the proximal and medial regions of the disc, where it is expressed at higher levels. To estimate the significance of this effect, we can consider Eq. (21): at constant [*dpp*] and [*tkv*], the steady-state concentration [*pMad*] is proportional to

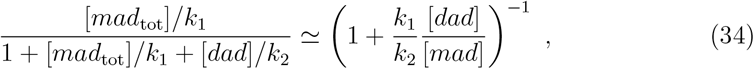

where we made the approximation, already discussed in Sec. 2.2.4, that the intracellular domain of Tkv is at all times either bound to Dad or Mad. Under wild-type conditions and in the proximal region, we have from Sec. 2.2.4 that 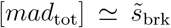 and from Sec. 2.2.2 that 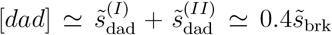, whence [*dad*]*/*[*mad*] ≃ 0.4, a factor of order unity. On the other hand, we expect *k*_2_*/k*_1_ ≪ 1, and indeed in Sec. 2.3 we saw that experimental gradients are most accurately recapitulated by the model when *k*_2_*/k*_1_ / 10^—2^. The factor in the right-hand side of Eq. (34) is thus expected to be small, of order 10^—1^ or smaller. Consequently, artificially setting [*dad*] = 0 under the same conditions should result in an increase in pMad signal by one order of magnitude or more.

This prediction can be tested against empirical evidence by considering discs where *dad* expression is inhibited, either everywhere by mutation, or in the dorsal compartment only by driving *dad*^*RNAi*^ with *apterous>*Gal4, see Fig. S10. Surprisingly, in contrast to our previous estimate, we observe only a mild increase in pMad signalling proximally (by approximately 30%) in both cases, with no qualitative changes in the shape of the Brk inverse gradient, except for a posterior shift. The limited enhancement of pMad signalling in the posterior compartment can be partly explained by noting that, as already discussed in Sec. 2.4.1.1, increased levels of pMad in the Dpp source downregulate morphogen production. Once again, this effect can be quantified by integrating the signal from GFP-Dpp over the source region, whence we estimate a 40% decrease with respect to wild-type production (Fig. S10I-J). The phosphorylation rate becoming limiting is unlikely as we observe no sign of saturation in the spatial pMad profile in either the *dad*^*RNAi*^ or *dad* mutant genotypes. Based on this evidence, we estimate that

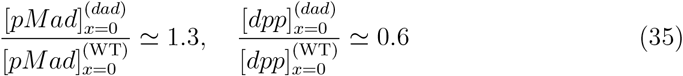

whereby, using Eq. (21), we conclude

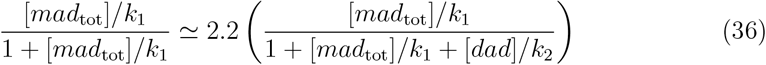

whence ([*dad*]*/k*_2_)(1 + [*mad*_tot_]*/k*_1_) ≃ 1.2. Using our estimates for [*dad*] and [*mad*_tot_] near the A-P boundary, we are left with two possibilities: the first requires 1 + [*mad*_tot_]*/k*_1_ ≃ 1, equivalently 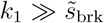, invalidating our assumption that the intracellular domain of Tkv is always occupied. In this case we find 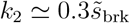 and the regime *k*_2_*/k*_1_ ≪ 1 remains accessible; alternatively, demanding 1 + [*mad*_tot_]*/k*_1_ ≃ [*mad*_tot_]*/k*_1_, which is in line with our assumption, we obtain *k*_2_*/k*_1_ ≃ 0.3, indicative of only a small difference in dissociation constants. The predicted gradients in the two regimes are shown in Fig. S10K-L. In either of these parameter regimes, the pMad and Brk gradients predicted by the model under wild-type, *brk* mutant or *dad* mutant conditions offer a noticeably poorer match to experimental data, which is recapitulated only at a qualitative level. A targeted dissection of the molecular mechanism underlying competitive inhibition by Dad might cast light on the issue of how excess proximal signalling is averted by the system, *e*.*g*. by identifying secondary low-affinity competitors of Mad (which would become important in the extreme case where Dad is completely removed) or by determining additional (feedback) mechanisms by which Dad inhibition is differentially controlled across the A-P axis. This second possibility seems consistent with the observation that pMad levels in wild-type and *dad* mutant conditions differ the most in the medial region (see Fig. S10E), while the wild-type Dad profile is monotonously decreasing with the distance from the A-P boundary.

## 3 Mathematical quantification of positional information: the role of noise

Within its range of validity, the model described above allows us to predict how the typical (i.e. average) concentration profile of different species of interest depends on the introduction or exclusion of certain regulatory elements, as well as changes in key parameters such as Dpp source production and diffusion, Mad phosphorylation rate by ligand-bound Tkv, expression levels, and so on. Accordingly, we concluded that, under wild-type conditions, Brk improves the readability of the Dpp morphogen gradient in the wing disc by decoding it into a smoother profile of *brk* expression. Indeed, one can roughly estimate the typical positional error associated with a cell’s estimate of its position based on the local properties of a (monotonic) morphogen gradient *s*(*x*) as

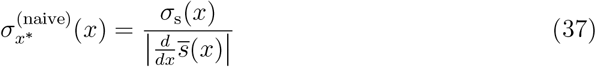

where *x* is the true position and *σ*_*s*_ is the position-dependent standard deviation associated with the variability (in time) of the morphogenetic signal *s* with average local concentration 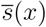 (*80*). We refer to this as a naive error, as it only relies on local information. It is clear from Eq. (37) that the error is large in regions where the signal gradient is flat (small *ds/dx*), whereby smoothing of an otherwise step-like gradient (compare the activity of the *brk* transcriptional reporter in *brk* mutant vs wild type) will naturally give a broader field of cells access to useful information about their position in the tissue (cf. expected *a posteriori* distribution in Fig. 4G-J). However, Eq. (37) also points to the importance of stochasticity, which we have hitherto neglected in our theoretical model. All the molecular mechanisms mentioned above (extracellular transport, binding, transcription and translation) are fundamentally stochastic in nature and contribute to different extents to temporal fluctuations in the concentration of any given species. A rigorous mathematical treatment of positional information at various levels of the Dpp signalling network would therefore require an extension of the model to explicitly include mechanisms, such as transcription, which have been coarse grained into effective rates in this work (see (*81*) for an example). Nevertheless, a rather general statement that can be made is that fluctuations in the concentration of a given species will decrease with increasing protein lifetime.

To see this, consider a protein *s* that is translated at a Poisson rate *τ* (*t*) fluctuating stochastically around a mean value 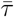 and degraded at rate *α*. In other words, the probability that a new protein is produced during the infinitesimal time window [*t, t* + Δ*t*] is *τ* (*t*)Δ*t*, taking *τ* to be independent of *s* for simplicity. The probability *P* (*n, t*) that there are *n* copies of *s* at time *t* evolves according to the master equation for a stochastic birth-death process

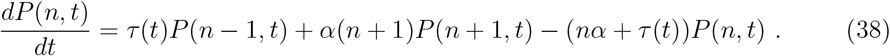

Accordingly, the first and second moment of the copy number for a particular realisation of the fluctuating production *τ* (*t*) satisfy

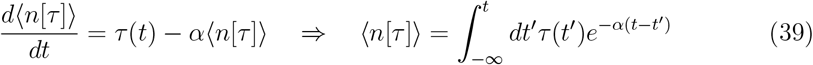

and

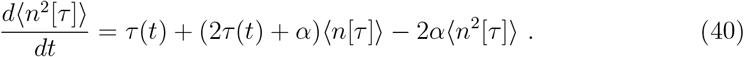

We now substitute (39) into (40) and perform an additional average, denoted ⟨⟨•⟩⟩, with respect to *τ* (*t*). At steady-state, we have that the mean and standard deviation of the copy number of *s* are given by

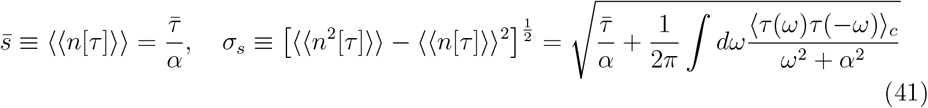

where 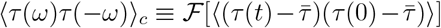 denotes the second stationary cumulant of *τ* in Fourier representation. Note that the production rate is a real number, *τ* ∈ ℝ, whence *τ* (*ω*)*τ* (−*ω*) = |*τ* (*ω*)|^2^ *>* 0 and thus the second contribution in the square root is non-negative: fluctuations in protein production (when independent of protein level itself) always increase copy number variability. In the absence of fluctuations in the translation rate, ⟨*τ* (*ω*)*τ* (−*ω*)⟩_*c*_ = 0, the copy number fluctuations are thus Poisson-like and scale like *α*^−1/2^, unlike the mean copy number which scales like *α*^−1^. In this regime, the naive error (37) can thus be made arbitrarily small by decreasing *α*, i.e. by increasing the protein lifetime. In the more general case where ⟨*τ* (*ω*)*τ* (−*ω*)⟩_*c*_ ≠0, we see that finite protein lifetime acts as a low pass filter, which buffers high frequency (fast) fluctuations in the translation rate, while leaving low frequency (slow) fluctuations unaffected. We argue that encoding positional information in the concentration gradient of Brk, which has a lifetime of approximately 10h, as opposed to that of pMad, which is dephosphorylated at a rate of about 0.07 min^—1^, thus offers an additional advantage in the form of time averaging. Note, however, that Brk being a more downstream product of the Dpp signalling pathway it is also more susceptible to intrinsic noise injected by upstream components, e.g. stochasticity in binding of pMad to the *brk* regulatory region.

**Figure S1:**
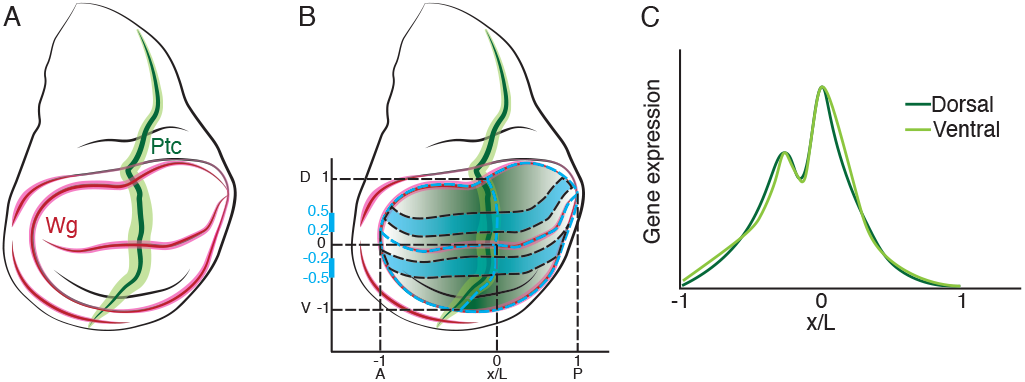
A coordinate system for segmentation. (**A**) Schematics showing the expression pattern of Wg (red) and Ptc (green) in a *Drosophila* wing imaginal disc. (**B**) Dashed blue lines mark the border of the wing disc pouch, as well as A-P and D-V boundaries. The origin of the coordinate system is set at the intersection between the A-P and D-V boundaries. The intersections between the A-P boundary and the pouch border are set to position -1 and 1 on the D-V axis. The intersections between the D-V boundary and the pouch border are set to position -1 and 1 on the A-P axis. The nuclei within 20%-50% along the D-V axis (indicated by blue lines) in the dorsal or ventral compartment were segmented for quantification, as illustrated in (C).

**Figure S2:**
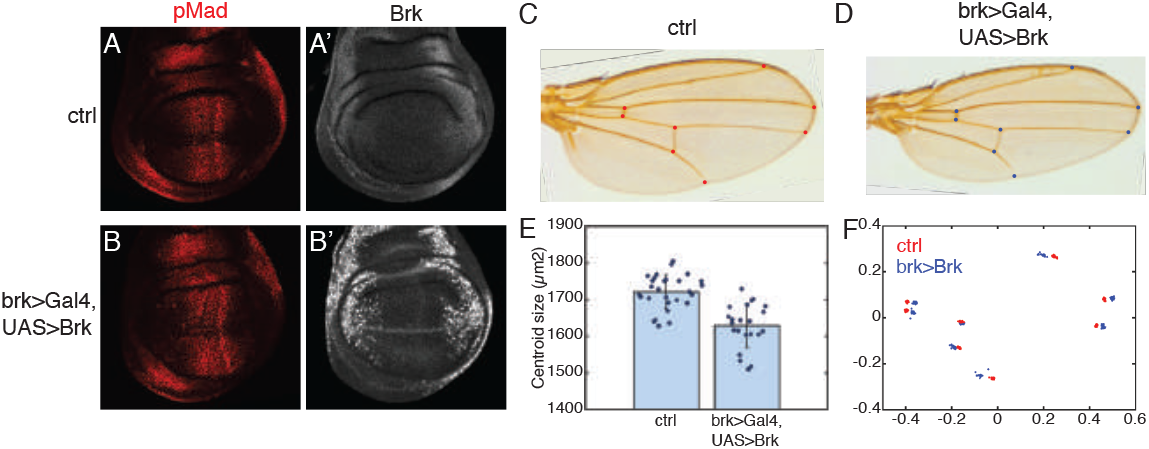
Effect of aberrant Brk expression on adult wing morphology. (**A**-**D**) pMad and Brk immunoreactivity in a wild type wing disc (ctrl) or a wing disc overexpressing Brk in the normal Brk-expression domain (brk*>*Gal4, UAS*>*Brk). Scale bar: 25 µm. Corresponding adult wings are shown in (C) and (D). Scale bar: 1 mm. Red and blue dots indicate vein landmarks used for quantitative analysis. (**E**) Mean centroid size and standard deviation calculated from segmented landmarks on wings of two genotypes. Dots mark values from individual wings. (**F**) Superimposed landmark positions of control (red) or Brk-overexpressing (blue) wings.

**Figure S3:**
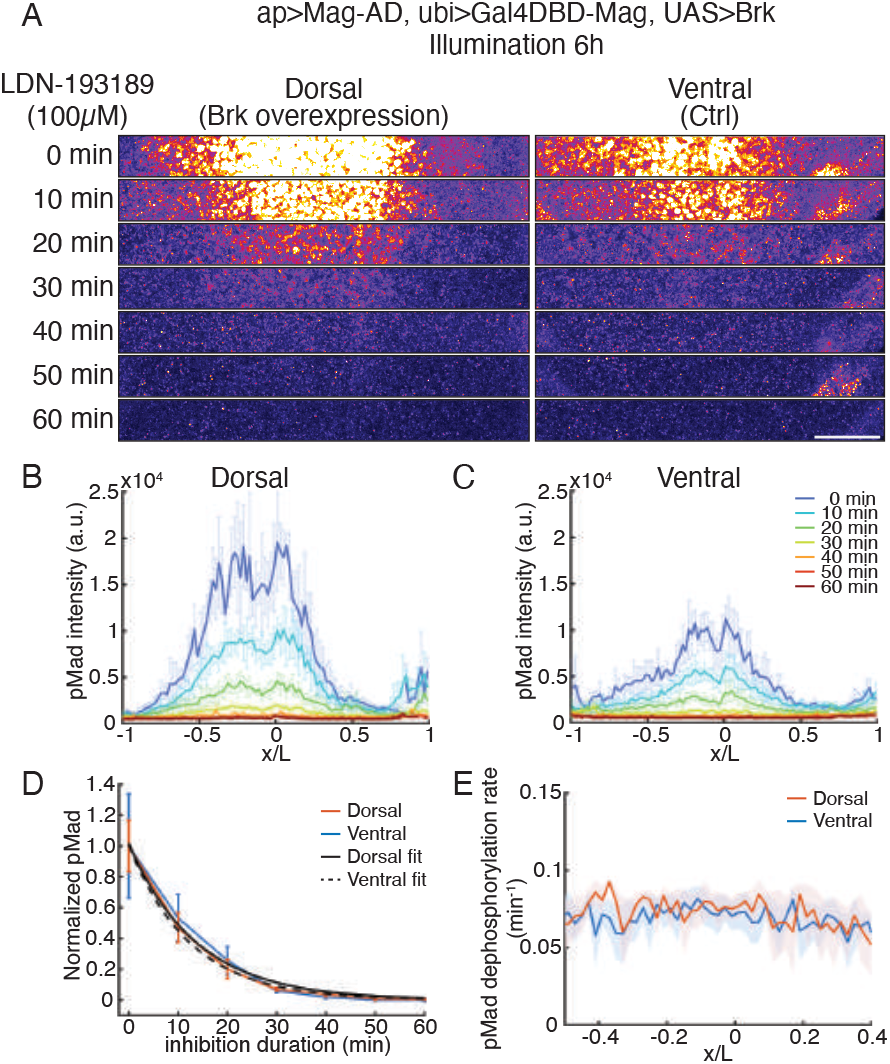
Brk does not modulate the rate of pMad’s dephosphorylation. (**A**) Brk overexpression leads to increased pMad immunoreactivity (warm color). Compare pMad in dorsal compartment (where Brk expression was induced by light for 6 hrs) and ventral compartment (control) at time 0 min. Treatment with LDN-193189 (BMP inhibitor) leads to progressive loss of pMad immunoreactivity, as shown in discs fixed and stained at regular intervals after drug treatment. Scale bar: 50 µm. (**B**-**C**) Spatial profiles of pMad intensity at different times after BMP inhibition. (**D**) Normalized pMad intensity quantified at 10% *x/L* (posterior peak) plotted against the duration of BMP inhibition. Exponential fits to the dorsal (solid lines) and ventral (dashed lines) compartment data are shown in black. (**E**) pMad dephosphorylation rate calculated from the fitted curves along the A-P axis in both compartments are the same, showing that Brk overexpression does not affect pMad’s dephosphorylation rate.

**Figure S4:**
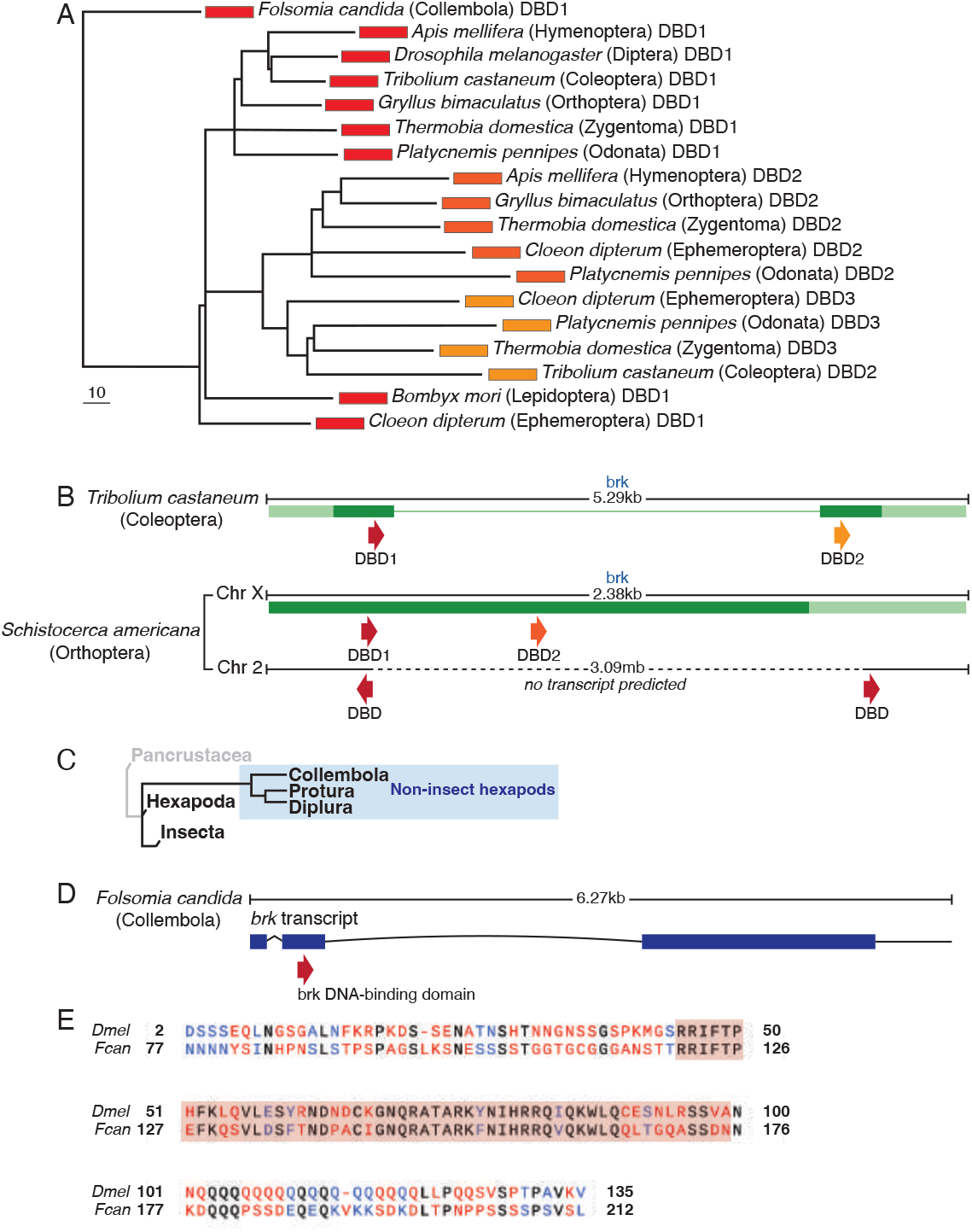
Conservation and divergence of the Brk protein among hexapods. (**A**) Neighbour-joining tree of Brk-DNA binding domains (DBDs) from representative hexapod species, based on BLOSUM62 distances. DBDs are color-coded as in Fig. 5A, with numbers showing their sequential positions in Brk proteins. Species order is shown in brackets. Scale bar represents 10 units of branch length. Notably, the second DBD of Coleoptera clusters with the third DBD of insects from basal groups (which contain three DBDs within the Brk protein), whereas the second DBD of Hymenoptera and Orthoptera clusters with the second DBD of these basal groups. This indicates that the triple DBD Brk is the ancestral form, and that derived forms have subsequently lost one or two DBDs. (**B**) Examples of alternative genomic organization of *brk* homologs. Top: In several Coleoptera species, including *Tribolium castaneum*, introns have evolved within the *brk* gene. Bottom: In certain Orthoptera species, such as *Schistocerca americana*, multiple Brk DBD homologs have been identified on different chromosomes. One chromosome harbours a transcript predicted to contain two DBDs, while another chromosome contains two single DBDs separated by a considerable chromosomal distance, without any predicted transcript within the region. (**C**) Phylogenetic diagram illustrating representative non-insect hexapod orders and their relationship with insects. (**D**) Gene structure of the putative *brk* homolog in *Folsomia candida*, a non-insect hexapod species. Exons are denoted by blue rectangles, and the conserved DBD is indicated by a red arrow. (**E**) Partial protein sequence alignment between *Drosophila* and *Folsomia* Brks, highlighting the DBD (shaded in red). Amino acid substitutions are denoted by red (different) and blue (similar) letters.

**Figure S5:**
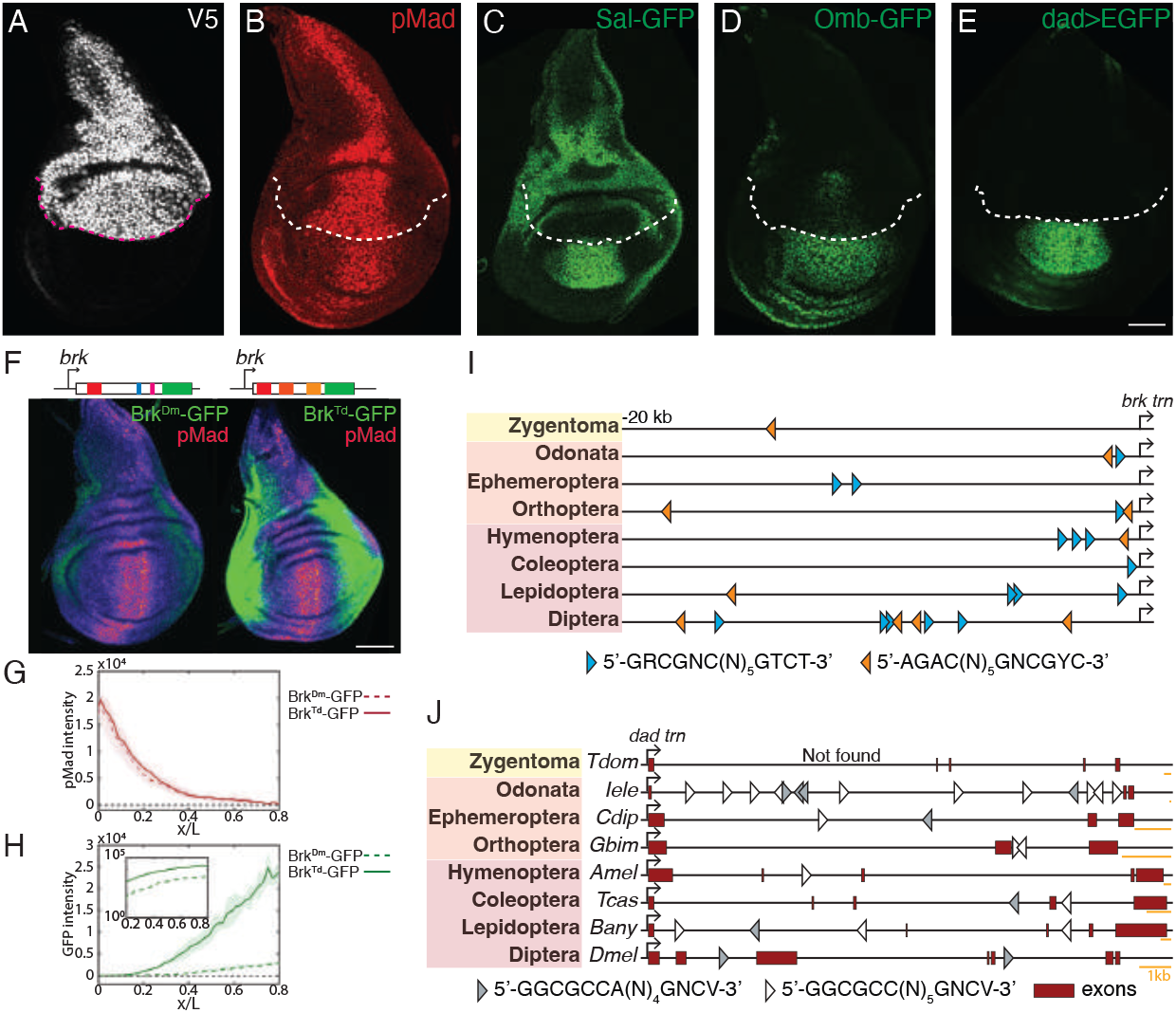
Brk protein but not its regulation is conserved across insect species. (**A**-**E**) Wing disc expressing V5-tagged *Thermobia* Brk in the dorsal compartment (*ap>Gal4, UAS>Brk*^*Td*^) stained for V5 (A) or pMad (B). GFP fluorescence in discs of the same genotype harbouring, in addition, Sal-GFP knock-in (C), Omb-GFP knock-in (D), or the transcriptional reporter dad*>*EGFP (E). Dashed lines indicate the D-V boundary. Scale bar: 50 µm. (**F**-**H**) Wing discs with endogenous *brk* coding region replaced with DNA encoding Brk^Dm^-GFP (left) or Brk^Td^-GFP (right), stained for pMad. Scale bar: 50 µm. Quantification of pMad (G) and Brk (H) intensities along the A-P axis is shown. Inset in (H) depicts the y-axis on a logarithmic scale, showing that Brk^Td^-GFP levels are approximately 10 times higher than Brk^Dm^-GFP levels at any A-P position. (**I**) Positions of putative pMad-Med-Shn-specific silencer motifs within the 20kb upstream of the *brk* gene coding region across representative insect orders. (**J**) Positions of putative Brk-binding sites within the introns regions of the *dad* genes across representative insect species.

**Figure S6:**
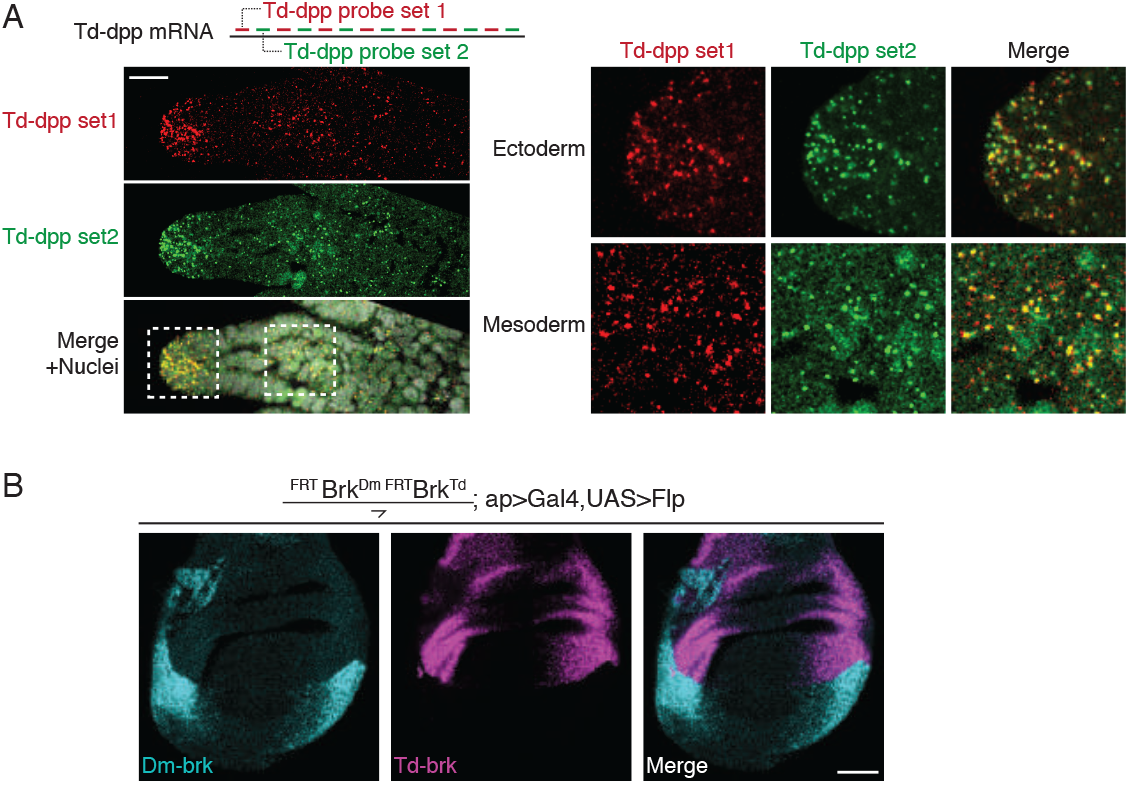
Validation of the *in situ* hybridization probes. (**A**) Dual-colour *in situ* hybridization of *dpp*^*Td*^ with two probe sets targeting alternating regions of the Td-dpp transcript. Scale bar: 25 µm.Areas of *Thermobia* limb primordia highlighted by dashed squares are shown at higher magnification on the right. Yellow dots in the merged images show the colocalization of the two probes, confirming their specificity. (**B**) *In situ* hybridization with *brk*^*Dm*^ and *brk*^*Td*^ of a wing disc expressing *Thermobia* Brk in the dorsal compartment and *Drosophila* Brk in the ventral compartment. Scale bar: 50 µm.

**Figure S7:**
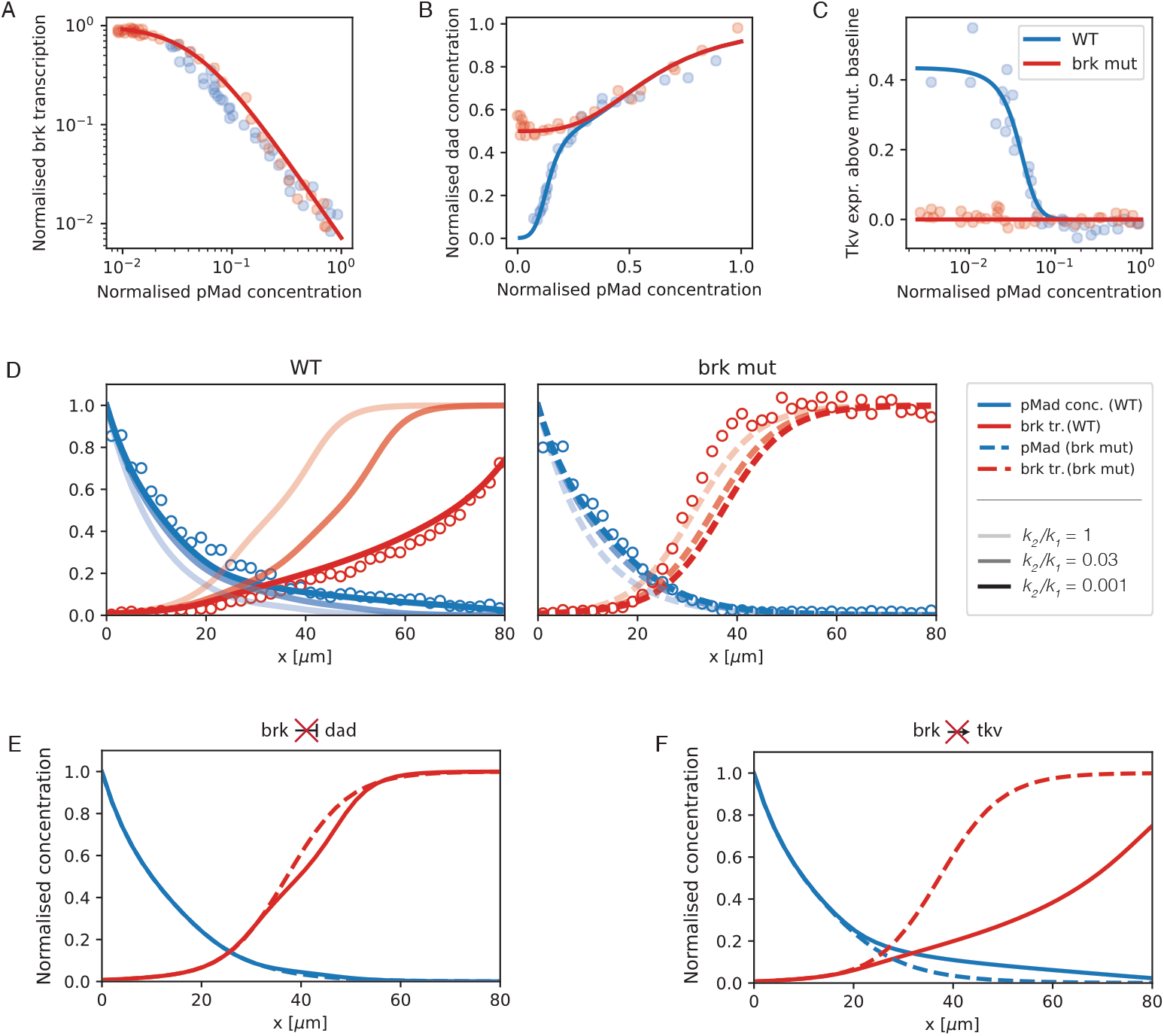
Formalisation of an experimentally informed mathematical model. (**A**-**C**) Input-output relationships between pMad and *brk* (A), *dad* (B), and *tkv* (C) in wild type (WT) and *brk* mutant backgrounds. Coloured circles represent experimental data points, while lines represent fitted Hill functions. Transcription of *brk, dad*, and *tkv* was measured using *brk>mCh, dad>lacZ*, and *tkv>lacZ* reporters, respectively. (**D**) Normalized spatial profiles of pMad immunoreactivity and *brk* transcription in WT and *brk* mutant backgrounds. Lines with different shading represent simulations with alternative *k*_2_*/k*_1_ ratios, and circles represent experimental data. (**E**-**F**) Simulated spatial profiles of pMad and *brk* transcription in a WT network (solid lines) compared to networks lacking the *dad* feedback branch (E) or the *tkv* feedback branch (F).

**Figure S8:**
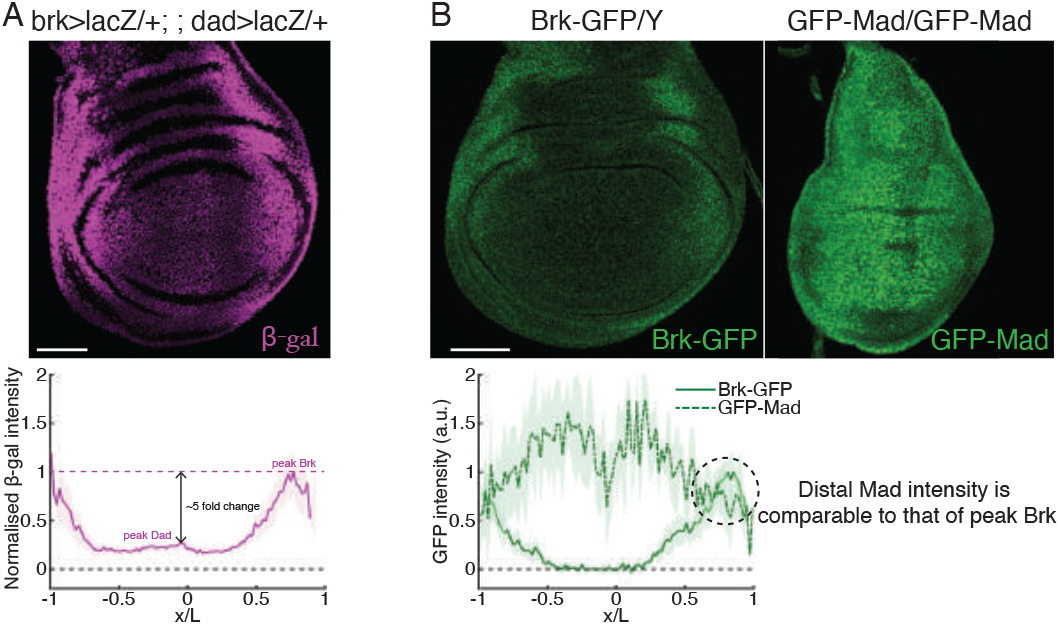
Model parametrisation using experimental measurements. (**A**) Activity of *brk>lacZ* and *dad>lacZ* enhancer traps in the same imaginal disc, as reported with anti-*β*-galactosidase (*β*-gal). The plot shows the mean nuclear intensity of *β*-gal against the A-P position. (**B**) GFP fluorescence from *brk-GFP* or *GFP-mad* knock-in alleles in imaginal discs imaged under identical conditions. The raw GFP intensity of both genotypes is plotted against the A-P position. Scale bars: 50 µm.

**Figure S9:**
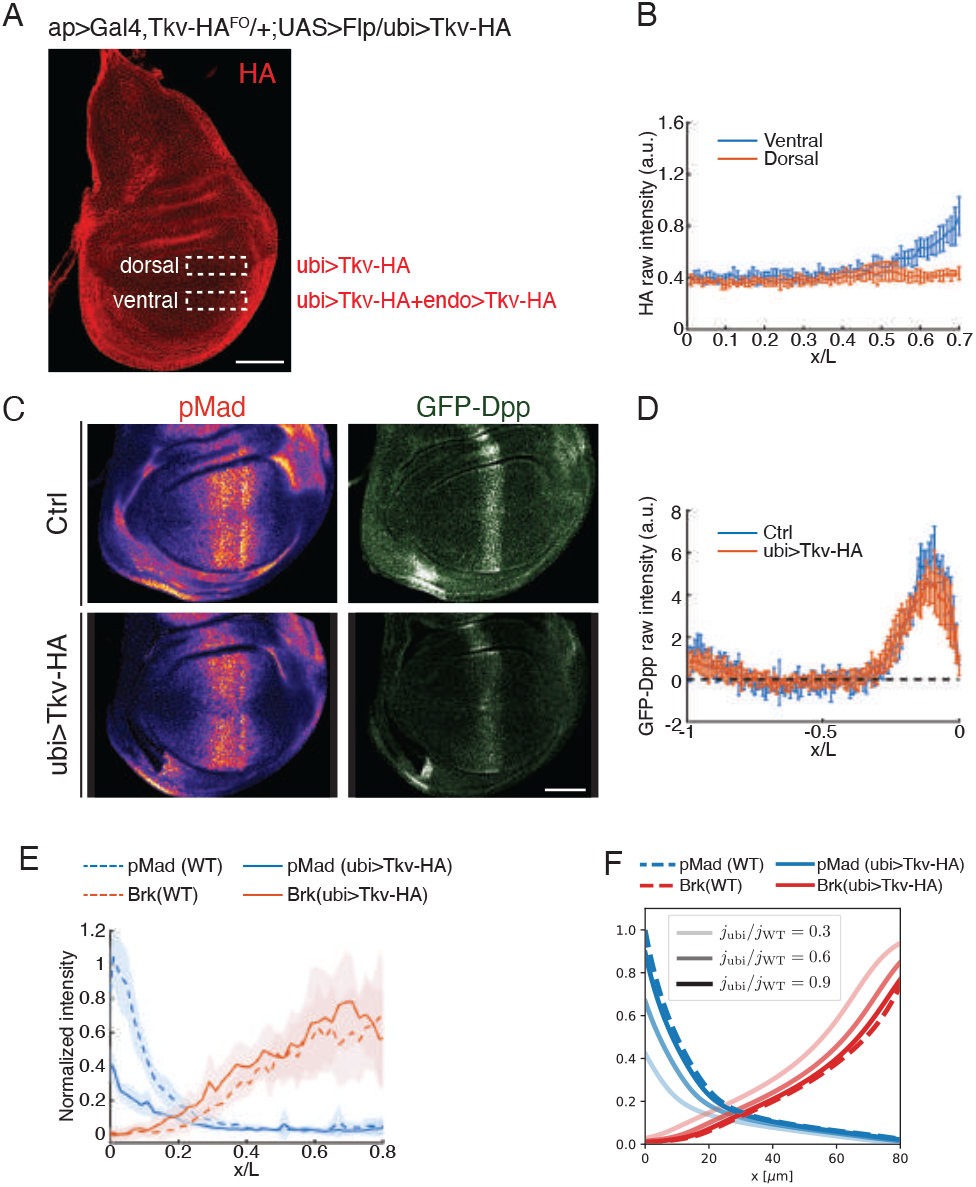
Establishment of signalling gradient with uniform receptor expression. (**A**) Comparing the production of Tkv from ubi*>*Tkv to that from the endogenous locus. In this genotype, only HA-tagged Tkv is produced, from a ubi*>*Tkv-HA transgene and from the endogenous locus (endo*>*Tkv-HA^FO^), which expresses Tkv-HA and can be inactivated by Flp. The endogenous locus was inactivated with ap*>*Gal4, UAS*>*Flp, leaving the *ubi>Tkv-HA* transgene as the only source of Tkv-HA in the dorsal compartment. Scale bar: 50 µm. (**B**) The spatial Tkv-HA profile is plotted in blue (ubi*>*Tkv-HA + endogenous Tkv-HA) and red (ubi*>*Tkv-HA). (**C**) Wing discs expressing GFP-Dpp from a knock-in allele in control (otherwise wild type) or ubi*>*Tkv-HA (lacking endogenous Tkv) discs. The samples were stained for pMad. Scale bar: 50 µm. (**D**) GFP-Dpp intensity along the A-P axis in the anterior compartment of control and ubi*>*Tkv-HA discs as in (C). (**E**-**F**) Spatial profiles of pMad and Brk in WT and ubi*>*Tkv-HA discs, derived from experimental measurements (E) or model simulation (F). Brk profiles were measured by fluorescence from a *brk-GFP* knock-in allele. Lines with different shades represent different values of the Dpp flux at the A-P boundary (compared to wildtype conditions), corresponding to different degrees of dpp updated in the producing stripe.

**Figure S10:**
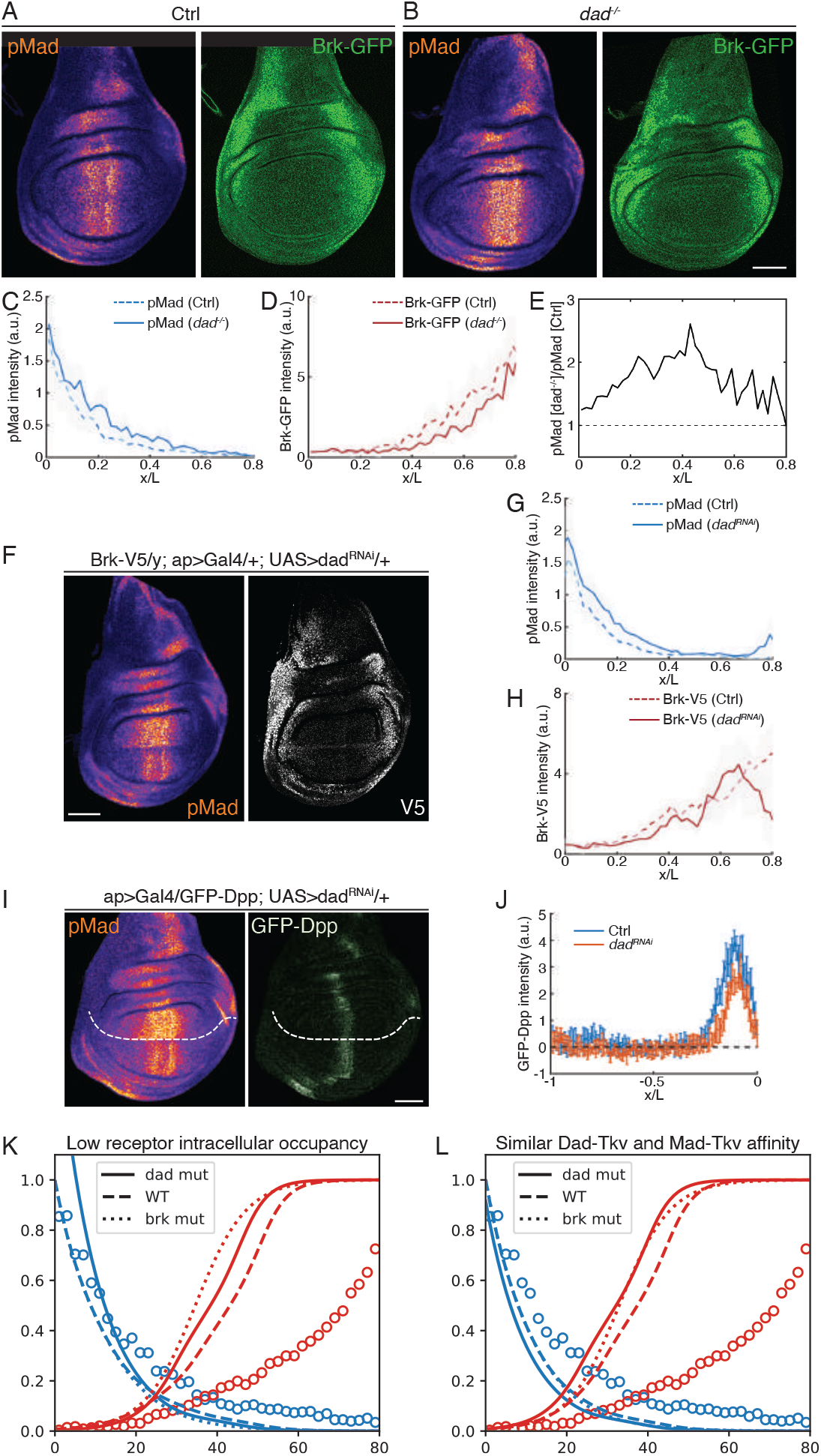
Establishment of signalling gradient in *dad* mutant. (**A**-**B**) pMad immunoreactivity and GFP fluorescence from a *brk-GFP* knock-in allele in control (*dad*^*+/+*^) and *dad* mutant (*dad*^*-/-*^) wing imaginal discs. (**C**-**D**) Intensity profiles of pMad and Brk-GFP in control (dashed lines) and *dad* mutant (solid lines) discs. (**E**) Quantification of the pMad [*dad*^*-/-*^]/pMad [WT] ratio along the A-P axis shows that the mid-axis region is most affected by the loss of Dad. (**F**) Effect of *dad* knockdown in the dorsal compartment (ventral compartment serves a control) on pMad and Brk expression (from a *brk-V5* knock-in). (**G**-**H**) Intensity of pMad and Brk-V5 in the control (dashed lines) and *dad*^*RNAi*^ (solid lines) compartments. (**I**) Effect of *dad* knockdown in the dorsal compartment (ventral compartment serves a control) on pMad and Dpp expression (from a *GFP-dpp* knock-in). (**J**) GFP-Dpp intensity in the anterior compartment of the control and the *dad*^*RNAi*^-expressing domains. (**K**-**L**) Simulated spatial profiles of pMad and Brk in WT, *dad* mutant and *brk* mutant conditions, testing two distinct parameter regimes. In (K), it is assumed that the intracellular domain of Tkv has a low occupancy by both Mad and Dad. In (L), it is assumed that the affinities for Dad-Tkv binding and Mad-Tkv binding are similar. Circles represent experiment data in WT. Scale bars: 50 µm.

**Table S1. (separate file): Brk protein sequence divergence across insect orders.** This table summarizes the divergence of Brk protein sequences across various insect orders, providing data on average protein lengths, DNA-binding domains, and interaction motifs (CtBP and Gro), along with sequence similarities and divergence.

**Table S2. (separate file): Silencer sites located within the 20 kb upstream region of *brk* genes across insect orders.** This table lists all identified pMad silencer sites located within the 20 kb upstream region of *brk* genes in various insect species. The sites are presented with their DNA sequence, distance from the *brk* transcription start site (TSS) or translation start site (if the 5’ UTR is unknown), and their 5’ or 3’ orientation.

**Table S3. (separate file): Brk-binding sites within the intronic region of *dad* genes across insect orders.** This table lists all identified Brk-binding sites located within the intronic region of *dad* genes in various insect species. The sites are presented with their DNA sequence, locations within the dad transcripts, and their 5’ or 3’ orientation.

**Table S4. (separate file): Genotypes analysed.** This table lists all the genotypes analysed in the corresponding figures.

**Table S5:** Fitted model parameters. All those parameters that do not appear explicitly in the table are set to unity without loss of generality by suitable choice of units.

| Parameter | Fitted value | Units |
| --- | --- | --- |
| $h_1$ | $1.60 \pm 0.04$ | - |
| $h_2$ | $3.0 \pm 0.3$ | - |
| $h_3$ | $3.1 \pm 0.4$ | - |
| $h_4$ | $5 \pm 1$ | - |
| $\sqrt{Dh/(r_{\text{dpp}}\tilde{s}_{\text{tkv}}^{(I)})}$ | 10 | $\mu m$ |
| $k_2/k_1$ | $10^{-3}$ | - |
| $k_3$ | $0.0459 \pm 0.0008$ | $[pMad]_{x=0}$ in wild-type |
| $k_4/\tilde{s}_{\text{brk}}$ | $0.150 \pm 0.004$ | - |
| $k_5$ | $0.60 \pm 0.02$ | $[pMad]_{x=0}$ in wild-type |
| $k_6/\tilde{s}_{\text{brk}}$ | $0.59 \pm 0.03$ | - |
| $\tilde{s}_{\text{dad}}^{(I)}/\tilde{s}_{\text{brk}}$ | 0.1 | - |
| $\tilde{s}_{\text{dad}}^{(I)}/\tilde{s}_{\text{dad}}^{(II)}$ | 1 | - |
| $\tilde{s}_{\text{tkv}}^{(I)}/\tilde{s}_{\text{tkv}}^{(II)}$ | 0.5 | - |
| $[mad_{\text{tot}}]/\tilde{s}_{\text{brk}}$ | 1 | - |

**Data S1 (separate file): Brk protein sequences across all analysed species**. This multiple sequence FASTA file contains all Brk protein sequences examined in this study. Each entry is labelled with a header that includes the insect order and species name.

## Notes

### Competing Interest Statement

The authors have declared no competing interest.

